# Single-cell imaging reveals spontaneous phenotypic sorting and bet-hedging in developing biofilms

**DOI:** 10.1101/2025.06.03.657734

**Authors:** Jung-Shen Benny Tai, Kee-Myoung Nam, Changhao Li, Japinder Nijjer, Sulin Zhang, Christopher M. Waters, Jing Yan

## Abstract

How phenotypic heterogeneity shapes biofilm architecture and development remains poorly understood. Motivated by this, we developed imaging tools to track intracellular levels of c-di-GMP, a key second messenger that controls the motile-to-sessile transition, in developing *Vibrio cholerae* biofilms at single-cell resolution. We show that c-di-GMP levels spontaneously bifurcate into a bimodal distribution that forms a spatially sorted pattern: High-c-di-GMP cells dominate the biofilm core while low-c-di-GMP cells localize to the periphery. Combining single-lineage tracing, mutant analysis, and agent-based modeling, we reveal that this pattern arises from differential viscosity and surface friction mediated by matrix-dependent interactions between cells and their microenvironments. We demonstrate that this heterogeneity and phenotypic sorting enable continuous emergence and shedding of planktonic cells, enhancing fitness in fluctuating environments. Our findings uncover a *differential drag mechanism* for pattern formation in multicellular systems, and expand the classical picture of biofilm lifecycle by highlighting the functional significance of phenotypic heterogeneity.

## Introduction

Biofilms are surface-attached, matrix-encased aggregates of bacterial cells that play critical roles in nature, health, and industry.^1^ The biofilm lifecycle starts with initial surface attachment, followed by extracellular matrix secretion and mass accumulation, and eventually dispersal and colonization of new niches.^2^ A key regulator of biofilm development is cyclic diguanylate (c-di-GMP), an intracellular second messenger that is synthesized and degraded by diguanylate cyclases (DGCs) and phosphodiesterases (PDEs), respectively (Figure 1A).^3^ In many bacterial species, c-di-GMP promotes the transition from a motile planktonic state to a sessile biofilm state by upregulating matrix production and downregulating motility-related processes, such as flagellar assembly.^4–6^ Beyond biofilm regulation, c-di-GMP also plays pivotal roles in regulating virulence, cell-cycle progression, and host-microbe symbiosis.^6–8^

**Fig. 1.**
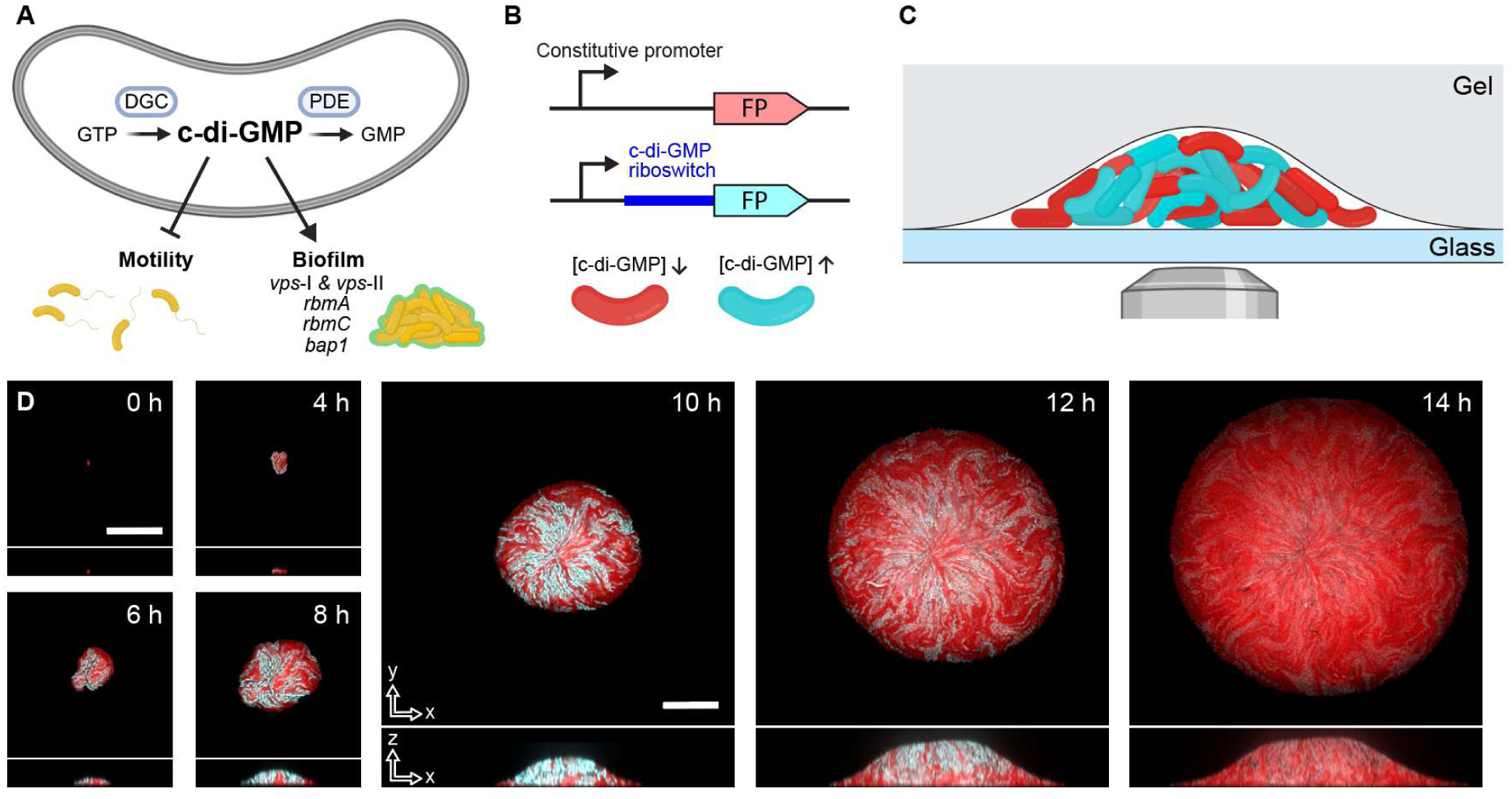
Single-cell imaging of *V. cholerae* biofilm development regulated by c-di-GMP. (**A**) Schematic of c-di-GMP metabolism by diguanylate cyclases (DGCs) and phosphodiesterases (PDEs) and its regulation of lifestyle switching in *V. cholerae*, by promoting matrix gene expression while suppressing motility. (**B**) A riboswitch-based fluorescent c-di-GMP biosensor was engineered into *V. cholerae* cells that constitutively express a red fluorescent protein (mScarlet-I). Cell appears red (cyan) when the intracellular c-di-GMP level is low (high) in overlaid images. (**C**) Schematic of a biofilm grown in the interstitial space between a glass substratum and an overlaid agarose hydrogel infused with growth medium and imaged through the glass from below. (**D**) Time-course confocal fluorescence images of both the constitutive marker (pseudo-colored red) and reporter (pseudo-colored cyan) channels of the bottom layer and vertical midplane cross-section of a WT *V. cholerae* biofilm with the c-di-GMP biosensor. Scale bars: 20 μm. (A), (B), and (C) were created using BioRender.com.

The classical model of the biofilm lifecycle suggests a coordinated process, in which cells synchronously transition between the planktonic and biofilm states. Indeed, collective processes such as quorum sensing are well-known to regulate biofilm formation,^9^ as do environmental conditions such as nutrient limitation.^10^ This generic picture, however, does not adequately account for the potential role of phenotypic heterogeneity in shaping biofilm development. Heterogeneity in gene expression is common in microbial communities, as has been made particularly evident in recent spatial transcriptomics studies,^11–13^ and is thought to confer fitness advantages through bet-hedging or division of labor.^14,15^ In biofilms, this heterogeneity is often attributed to local chemical cues received by cells at different locations, arising from gradients of nutrients and signaling molecules.^16,17^ However, little is known about the spatiotemporal dynamics of heterogeneity during biofilm formation at the level of individual cells, largely due to challenges in tracing lineages and single-cell phenotypes in compact, three-dimensional (3D) biofilms. As such, despite its widely acknowledged importance in facilitating pathogenicity and elevated antibiotic tolerance in biofilm-associated infections,^18,19^ how phenotypic heterogeneity shapes biofilm architecture and development has remained largely mysterious.

Here, by developing new imaging tools to track individual cells, single lineages, and their intracellular [c-di-GMP] in *Vibrio cholerae* biofilms, we uncover extensive heterogeneity in c-di-GMP signaling, which manifests as a bimodal [c-di-GMP] distribution across a growing biofilm. Moreover, these phenotypically heterogeneous biofilms spontaneously and robustly develop a spatially sorted pattern, in which high-c-di-GMP cells dominate the biofilm core and low-c-di-GMP cells localize to the periphery. Combining experiments and agent-based modeling, we demonstrate that this sorting arises from differential interactions between low-and high-c-di-GMP cells and their respective microenvironments, in a manner superficially analogous to but mechanistically distinct from cell sorting in eukaryotic systems.^20–23^ In contrast with the conventional view of phenotypic heterogeneity as a consequence of position-dependent chemical environments,^16,17^ we show that spontaneous phenotypic heterogeneity within a growing biofilm can also dictate the positional fates of the biofilm-dwelling cells. Finally, we show that phenotypic heterogeneity and sorting facilitate continuous shedding of planktonic, low-c-di-GMP cells from a biofilm, which enhances fitness and colonization efficiency in fluctuating environments. Our findings expand upon the classical view of the biofilm lifecycle and demonstrate a novel mechanism of pattern formation in multicellular systems.

## Results

### Single-cell imaging enables quantification of c-di-GMP levels in biofilm cells

To quantify the intracellular [c-di-GMP] of individual cells in growing *V. cholerae* biofilms, we engineered a riboswitch-based biosensor^24^ into wild-type (WT) *V. cholerae* cells to express a fluorescent protein (FP, mNeonGreen or SCFP3A depending on the experiment) in a c-di-GMP-dependent manner, based on binding between c-di-GMP and an upstream riboswitch (Figures 1B, S1A, and S1B). The cells also constitutively express a red FP marker (mScarlet-I) for cell segmentation and calibration of the c-di-GMP biosensor signal. Biofilms were grown in the interstitial space between a glass substratum and an overlaid agarose hydrogel, such that all cells descending from the founding cell remain within the imaging field (Figures 1C, S1C, and S1D). Biofilms were imaged using a confocal microscope in both the marker and reporter channels at 1-h intervals (Figure 1D; Video S1) throughout the entire developmental process. Surprisingly, we found strong heterogeneity in c-di-GMP biosensor fluorescence across the cells in each biofilm; moreover, the biofilms exhibited a spatial pattern, in which high- and low-c-di-GMP cells within the bottom layer of each biofilm spontaneously segregated to the center and periphery, respectively. We also observed that cells close to the center align preferentially in the radial direction;^25–27^ in contrast, cells in the periphery were disordered.

We segmented the growing biofilms at each timepoint into individual cells using a custom algorithm (Figure 2A).^27,28^ The c-di-GMP fluorescence intensity (*I*_c-di-GMP_) of each cell was quantified using the reporter fluorescence, normalized by the constitutive marker fluorescence and further calibrated for the change in fluorescence signal with distance from the glass substratum (Figures S1E and S1F). Quantification of *I*_c-di-GMP_ showed that, on average, [c-di-GMP] starts at a low value and increases as the biofilm develops and peaks at 9–10 h, before eventually decreasing to a low value by 17 h (Figure 2B). Remarkably, significant heterogeneity in [c-di-GMP] arises spontaneously during exponential phase, with a marked bimodal distribution fully developed after 8 h. Consistent with the visual inspection, the spatiotemporal evolution of *I*_c-di-GMP_ shows that high-c-di-GMP cells are enriched close to the center of the bottom layer in each biofilm (Figure 2C).

**Fig. 2.**
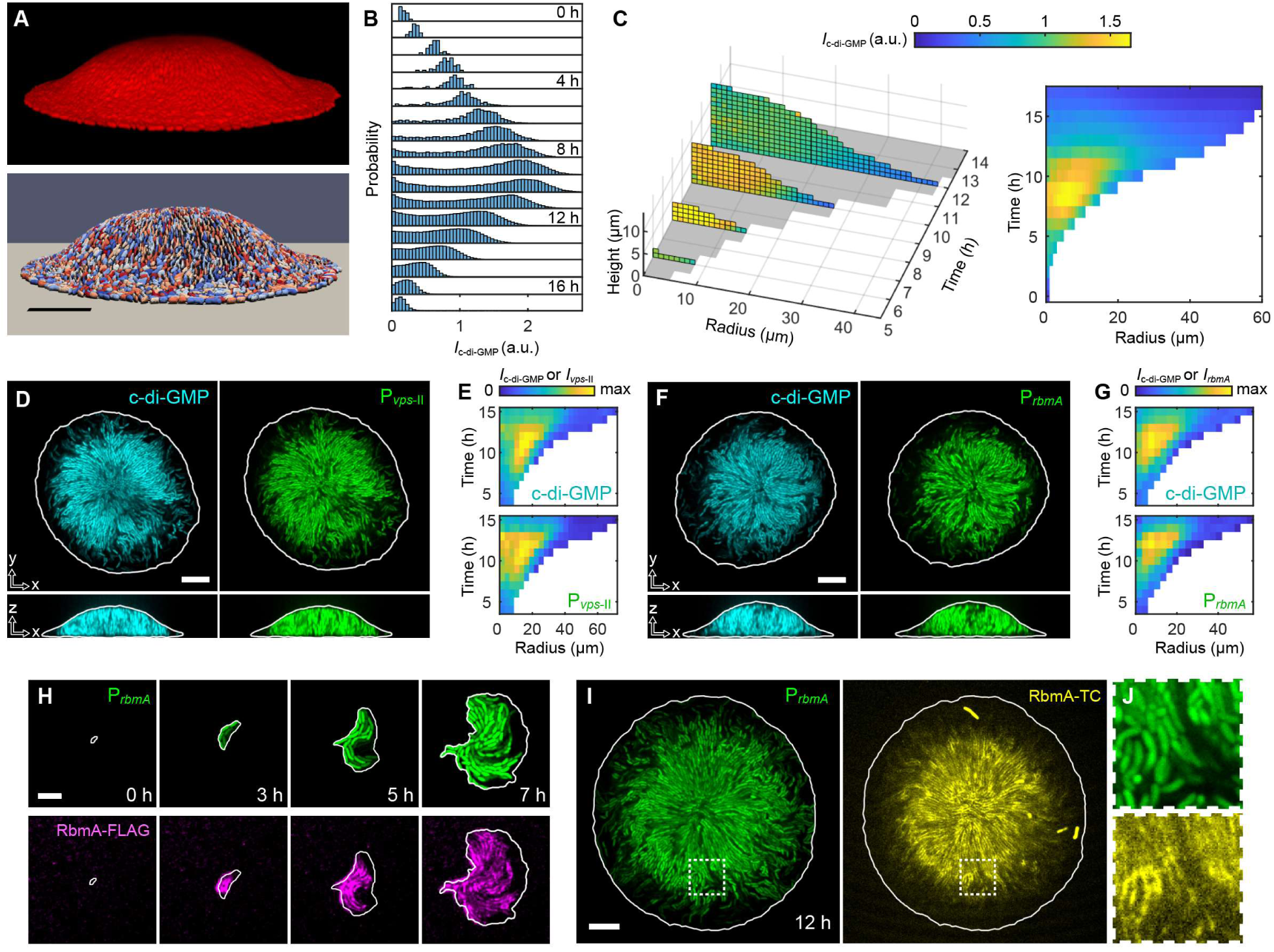
Single-cell quantification of c-di-GMP and matrix gene expression and localization. (**A**) 3D volume rendered from confocal images (*top*) of a WT *V. cholerae* biofilm at 11 h and its segmentation into 5028 individual cells (colored randomly, *bottom*). (**B**) Distribution of *I*c-di-GMP across cells within WT *V. cholerae* biofilms over time. (**C**) Spatiotemporal evolution of azimuthally averaged *I*c-di-GMP, shown as height vs. radius heatmaps at different timepoints (*left*) and a time vs. radius heatmap for the bottom layer (*right*) of WT biofilms. Data in (B, C) were obtained from 11 biofilms from 3 independent experiments. (**D**) Cross-sectional images of a biofilm grown for 11 h from cells simultaneously expressing the c-di-GMP biosensor (*left*) and the *vps*-II transcriptional reporter (*right*). (**E**) Time vs. radius heatmaps of c-di-GMP level and *vps*-II expression in cells in the bottom layer of the biofilm shown in (D). (**F**) Cross-sectional images of a biofilm grown for 12 h from cells simultaneously expressing the c-di-GMP biosensor (*left*) and the *rbmA* transcriptional reporter (*right*). (**G**) Time vs. radius heatmaps of c-di-GMP level and *rbmA* expression in cells in the bottom layer of the biofilm shown in (F). (**H**) P*rbmA* activity (*top*) in individual cells and the corresponding immunofluorescence staining of RbmA-3×FLAG in the extracellular space (*bottom*) in an early biofilm. (**I**) P*rbmA* activity (*left*) and the corresponding staining of tetracysteine (TC)-tagged RbmA with a biarsenical dye (ReAsH-EDT2; *right*) in a mature biofilm. Some dead cells are seen as bright cells in the TC channel. (**J**) Zoomed-in view of the dashed boxes in (I), showing the correlation between heterogeneity in P*rbmA* activity and RbmA staining. White lines in (D, F, H, I) delineate biofilm boundaries based on the constitutive marker. Scale bars: 10 μm in (A, D, F, I) and 5 μm in (H).

To demonstrate the correlation between heterogeneity in [c-di-GMP] and the expression levels of biofilm matrix components, we engineered dual reporter strains, each containing the c-di-GMP biosensor and a transcriptional reporter for one of the matrix genes (Figure S1A). The major components of the *V. cholerae* biofilm matrix are <u>V</u>ibrio <u>p</u>oly<u>s</u>accharide (VPS)^29^ and three adhesion proteins: RbmA,^30,31^ which facilitates cell-cell adhesion, and Bap1 and RbmC,^32–34^ which facilitate cell-surface adhesion. The transcription of *vps*-II, the second operon of VPS biogenesis genes, and *rbmA* both show similar levels of heterogeneity to the c-di-GMP biosensor, and strong spatiotemporal correlation with *I*_c-di-GMP_ at the single-cell level (Figures 2D–2G). This strong correlation is consistent with known pathways in which c-di-GMP upregulates the expression of *vps* operons and genes encoding the matrix proteins via the transcription factors VpsR and VpsT.^5^ Moreover, the low *vps* expression level at the periphery is consistent with the locally disordered cell organization, since matrix production has been shown to be required for radial ordering.^27^ Furthermore, by labeling RbmA with either a 3×FLAG^35^ or a tetracysteine (TC) tag^36^ and visualizing with immunofluorescence or a biarsenical dye, respectively, we found that RbmA molecules in biofilms closely associate with cells with high P*_rbmA_* activity and [c-di-GMP] throughout biofilm growth (Figures 2H–2J, S1G, and S1H). Overall, our data reveal substantial heterogeneity and spatiotemporal sorting in c-di-GMP level within growing *V. cholerae* biofilms, which in turn give rise to heterogeneity and sorting in downstream production and localization of the biofilm matrix.

### Matrix composition impacts c-di-GMP pattern and cellular alignment

To identify mechanisms that could give rise to the observed c-di-GMP pattern, we quantified the spatiotemporal evolution of *I*_c-di-GMP_ and cellular organization in biofilms formed by various matrix gene deletion mutants (Figures 3A–3C). To quantify the “sortedness” of the c-di-GMP pattern, we calculated Spearman’s rank correlation coefficient (ρ_s_) between *I*_c-di-GMP_ and the radial position of each cell. According to this definition, a monotonically decreasing c-di-GMP level from center to periphery corresponds to ρ_s_ = −1 and a monotonically increasing c-di-GMP level corresponds to ρ_s_ = 1, while ρ_s_ = 0 suggests an unsorted pattern (Figure 3D). We also quantified the radial alignment of each cell by *S* ≡ 2(*n^^^*_ǁ_ ⋅ *r^^^*)^2^ − 1, where *n^^^*_ǁ_ is the in-plane orientation of the cell an *r^^^* is the unit vector from the origin of the biofilm to the cell center, so that *S* = 1 and –1 correspond to perfect radial and tangential cell alignment, respectively (Figure 3D).

**Fig. 3.**
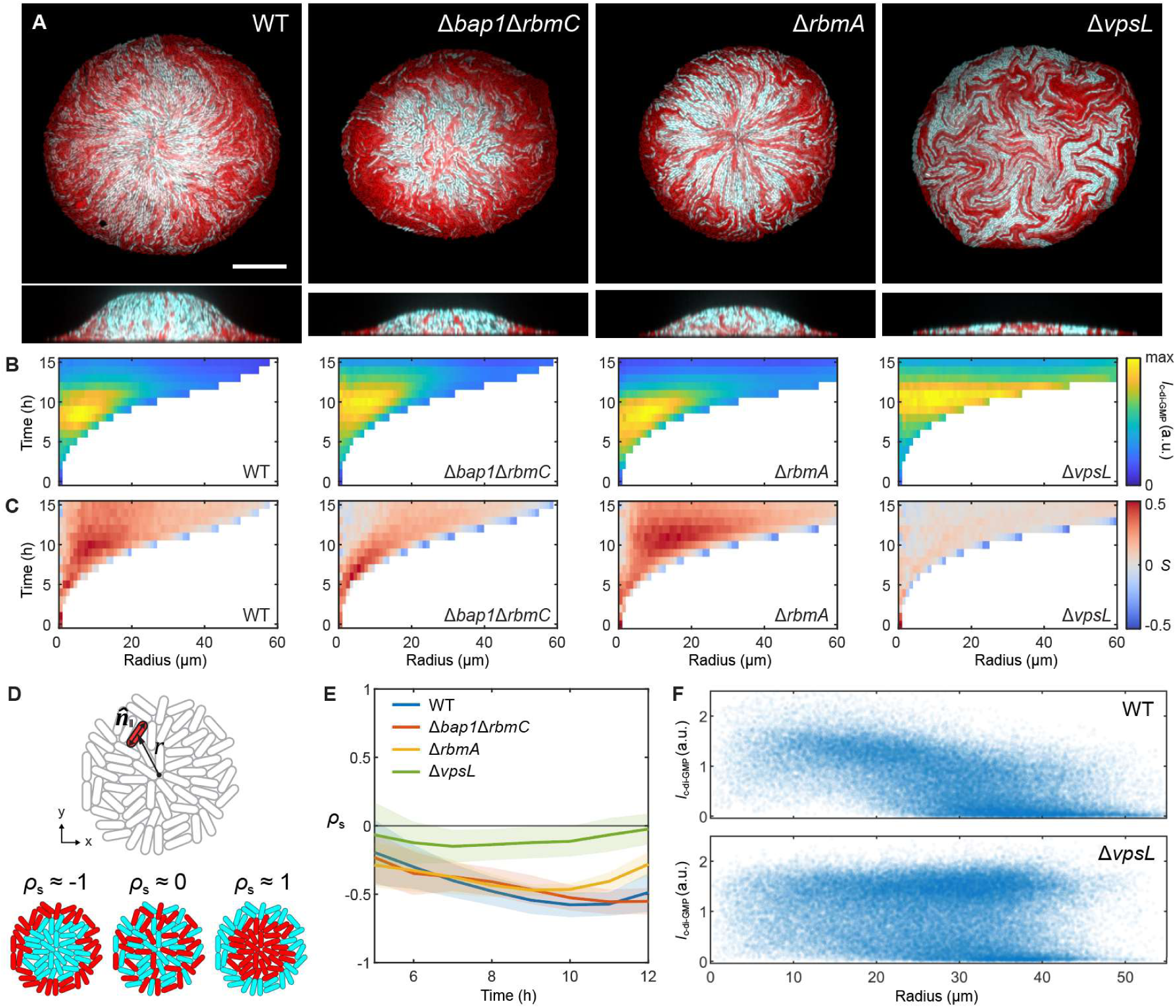
Matrix components impact phenotypic sorting in *V. cholerae* biofilms. (**A**) Bottom layers (*top*) and vertical midplane cross-sections (*bottom*) of biofilms formed by WT, Δ*bap1*Δ*rbmC*, Δ*rbmA*, and Δ*vpsL V. cholerae* strains (left to right). Scale bar: 20 μm. (**B**) Time vs. radius heatmaps of locally averaged *I*c-di-GMP of cells in the bottom layer of WT and mutant biofilms. (**C**) Time vs. radius heatmaps of locally averaged radial alignment, *S* ≡ 2(*n^^^*_ǁ_ ⋅ *r^^^*)^2^ − 1, of cells in the bottom layer of WT and mutant biofilms. (**D**) *Top*: Schematic of in-plane orientation, *n^^^*_ǁ_, of an individual cell and its distance, *r*, to the origin of the biofilm. *Bottom*: Illustrations of example patterns of c-di-GMP phenotypes and their corresponding Spearman’s rank correlation coefficients (*ρs*) between c-di-GMP level and *r*. (**E**) *ρs* vs. time for WT and mutant biofilms. Solid lines and shading indicate mean ± s.d. (**F**) Scatterplots of *I*c-di-GMP vs. radius of cells in the bottom layer of WT and Δ*vpsL* biofilms. The plots reflect, for each biofilm, the timepoint at which the bottom layer contained ∼5000 cells (radii of biofilms were ∼40 μm). Data in (A, B, C, E, F) were obtained from 11–25 biofilms from 3 independent experiments for each strain.

In WT biofilms, ρ_s_ shows a downward trend starting from 5 h (before which quantification is unreliable due to the small number of cells) and reaches a minimum value of ρ-_s_ = −0.58 at 10 h (Figures 3E and S2A). In Δ*bap1*Δ*rbmC* biofilms, wherein the two biofilm-specific surface adhesins are absent,^32–34^ the sorted c-di-GMP pattern remains and ρ_s_ is not significantly different from that of WT biofilms. In Δ*rbmA* biofilms, sortedness is slightly tempered (minimum ρ-_s_ = −0.47 at 9 h). Remarkably, in colonies formed by Δ*vpsL* cells that do not produce VPS (so that RbmA, Bap1, and RbmC, whose functions depend on VPS binding, are all nonfunctional),^29,31,34^ the sorted c-di-GMP pattern is largely abolished (minimum ρ-_s_ = −0.15 at 7 h). These data suggest that either cell-cell or cell-surface adhesion is sufficient to generate the sorted pattern, whereas removing both eliminates it. Unlike the c-di-GMP pattern, the radial ordering depends critically on Bap1/RbmC: Biofilms with these two proteins present (WT and Δ*rbmA*) exhibit much higher radial ordering than those without them (Δ*bap1*Δ*rbmC* and Δ*rbmA*Δ*bap1*Δ*rbmC*, Figures 3C, S2B, and S2C). Moreover, radial ordering is completely lost in Δ*vpsL* colonies—this is consistent with previous studies showing that Bap1/RbmC/VPS-mediated cell-surface adhesion is essential for initiating the reorientation cascade that leads to radial alignment.^27^ The different genetic requirements for radial ordering and c-di-GMP-based sorting also suggest that they are not causally related. Notably, we found that the non-motile Δ*pomA* mutant strain also exhibits sorting (Figure S2D), which implies that sorting does not depend on the motility of low-c-di-GMP cells, but rather on the composition of the matrix surrounding the cells.

To further analyze the sorting pattern, we agglomerated all data from independent experiments pertaining to each strain and plotted the distribution of *I*_c-di-GMP_ vs. radial position of cells in the bottom layer of biofilms with radii ∼ 40 μm (around 12 h; Figures 3F and S2E). For Δ*vpsL* colonies, this plot shows two horizontal bands covering the entire range of radial positions, corresponding to the low- and high-c-di-GMP cells. In contrast, in WT and Δ*bap1*Δ*rbmC* biofilms, the high-c-di-GMP band dominates the core while the low-c-di-GMP band dominates the periphery, resulting in a significantly nonzero value of ρ_s_. Δ*rbmA* biofilms show a similar enrichment of low-c-di-GMP cells at the periphery, but the high-c-di-GMP cells are more diffusely distributed across the entire range of radial positions. Collectively, the various c-di-GMP patterns formed by these matrix mutants suggest that matrix production plays a significant role in shaping the c-di-GMP pattern, and different matrix components may contribute differently to sorting, based on the physical interactions they confer.

### Single-lineage tracing reveals c-di-GMP-dependent cell dynamics

To uncover the mechanisms underlying the observed sorting, we investigated how [c-di-GMP] is associated with the dynamics of cells along individual lineages within a biofilm. To do this, we developed a new single-lineage tracing technique that takes advantage of an intracellular fluorescent punctum assembled from constitutively expressed mNeonGreen (mNG)-μNS^28,37,38^ and an asynchronous imaging protocol (Methods). By imaging the puncta at high time resolution (every 3–5 min) and the cells at a lower time resolution (every 30 min) and associating each punctum trajectory with a single lineage of cells, we mitigated photobleaching and phototoxicity while enabling simultaneous tracing of individual lineages within the 3D biofilms and tracking their *I*_c-di-GMP_ (Video S2). We found that trajectories that extend to the expansion front are dominated by low-c-di-GMP cells, while high-c-di-GMP cells follow trajectories that are more localized at the center (Figures 4A and S3A). Most interestingly, many lineages exhibit fluctuations in *I*_c-di-GMP_ over the course of biofilm growth, with variable “lifetimes” in the low- and high-c-di-GMP states (Figures 4B and S4). This suggests that c-di-GMP heterogeneity in biofilms at the community level is a consequence of phenotypic fluctuations at the single-cell level.

**Fig. 4.**
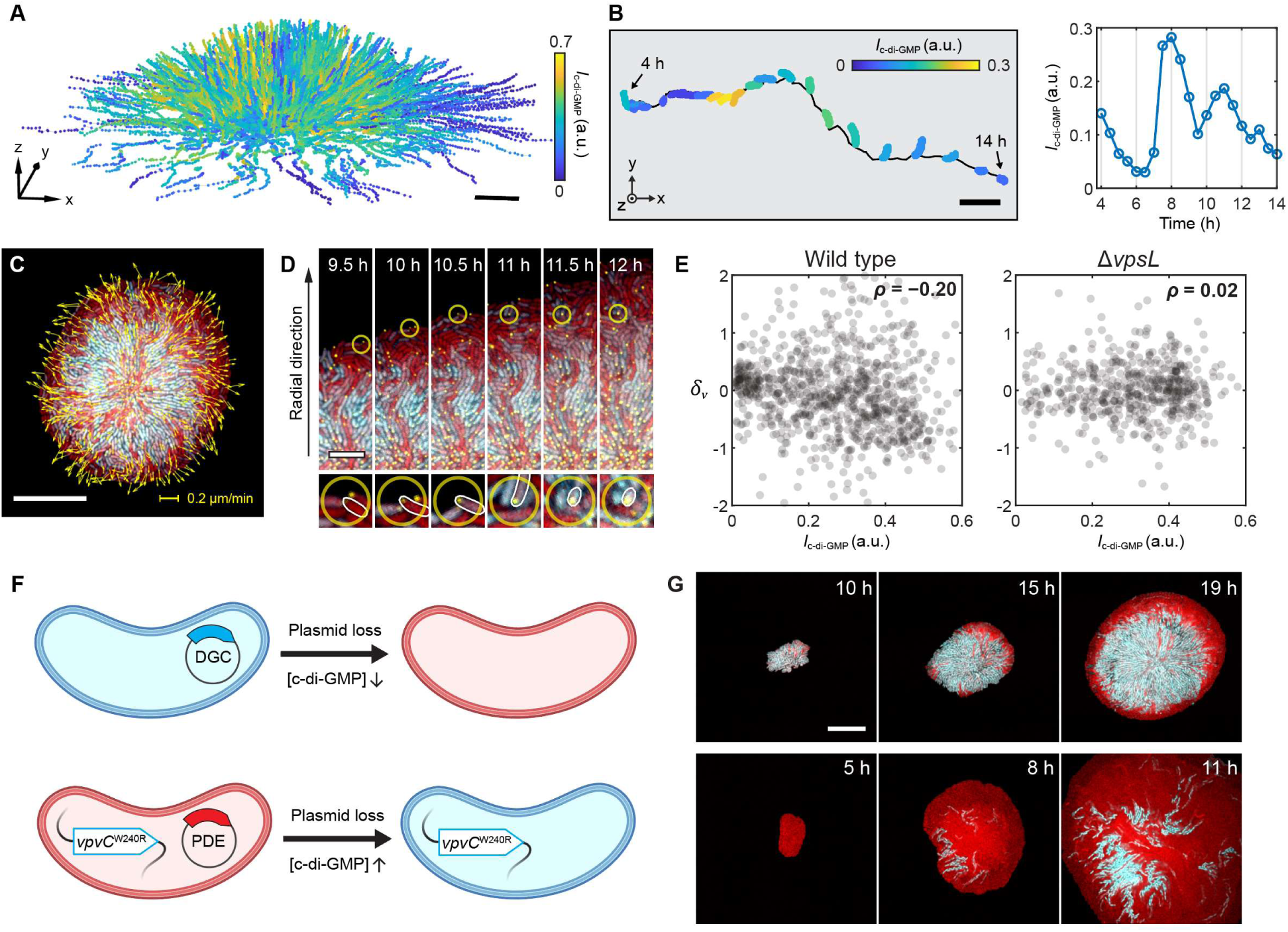
Single-lineage trajectories and engineered binary populations reveal c-di-GMP-dependent cell dynamics. **(A)** Single-lineage trajectories in a *V. cholerae* biofilm expressing mNG-μNS puncta, color-coded by instantaneous *I*c-di-GMP, between 0 and 12 h. The 1000 trajectories with the longest tracking durations are shown. (**B**) An example single-lineage trajectory as well as segmented cells in the same biofilm in (A) from 4 to 14 h (*left*), showing fluctuations in *I*c-di-GMP (*right*). (**C**) Confocal image of the bottom layer of the same biofilm in (A) at 11 h, showing both constitutive (red) and c-di-GMP biosensor (cyan) fluorescence, as well as instantaneous in-plane velocities (yellow arrows) tracked with mNG-μNS puncta (yellow). (**D**) Snapshots showing the cells along a single lineage (highlighted by yellow circle) with increasing *I*c-di-GMP and the corresponding self-arrest in its outward motion. *Bottom*: zoomed-in view of the cells. (**E**) Spearman’s rank correlation between δ_v_ and *I*c-di-GMP for cells in the bottom layer of a WT (*left*) and Δ*vpsL* (*right*) biofilm, both at 11 h. δ_v_ ≡ (*v*_r_ − *v*^-^_r_(*r*))/*v*^-^_r_(*r*), where *v*^-^_r_(*r*) is the average radial velocity at radius *r* from the origin of the biofilm. (**F**) Schematics of binary populations generated by the stochastic loss of a plasmid encoding a DGC in a low-c-di-GMP background (*top*) or a plasmid encoding a PDE in a high-c-di-GMP background (*bottom*). (**G**) Snapshots of the bottom layer of a biofilm in which individual cells irreversibly switch from high-to low-c-di-GMP (*top*) or *vice versa* (*bottom*) through stochastic plasmid loss. Scale bars: 5 μm in (A, B, D) and 20 μm in (C, G). (F) was created using BioRender.com.

We also found that, in some lineages, cells exhibit a strong anti-correlation between their *I*_c-di-GMP_ and instantaneous velocity (Figures 4C and 4D), which, in the biofilm context, arises not from motility but from the growth-induced flow that carries all cells radially outward (Figure S2D). This anti-correlation can be observed in an example single-lineage trajectory (Figures 4D and S3C; Video S3), in which a cell at the periphery exhibited low *I*_c-di-GMP_ at 9.5 h, then, within 1.5 h, the cell bearing the same punctum increased its *I*_c-di-GMP_ while concomitantly decreasing its velocity, and eventually became stationary at 11 h. We also observed the inverse scenario, in which a decrease in *I*_c-di-GMP_ was accompanied by the inception of outward motion in an initially stationary cell (Figures S3D and S3E, Video S3). To establish a statistical correlation between *I*_c-di-GMP_ and the outward motion of cells, we compared *I*_c-di-GMP_ with δ_v_, the difference between the radial velocity (*v*_r_) of each cell and the average *v*_r_ at a given radius (Methods). In WT biofilms, we found that δ_v_ is negatively correlated with *I*_c-di-GMP_ (ρ = −0.20; Figures 4E and S3F), meaning that a low-c-di-GMP (high-c-di-GMP) cell tends to move faster (more slowly) than the average cell at the same radial position. In Δ*vpsL* colonies, no significant correlation between δ_v_ and *I*_c-di-GMP_ was observed. Finally, we found that *I*_c-di-GMP_ also varies significantly in the circumferential direction, mainly due to radially segregated “lanes” consisting of cells from closely related lineages of either low- or high-c-di-GMP cells (Figure 4C). Consequently, the variations in *I*_c-di-GMP_ and *v*_r_ along the circumferential direction are also negatively correlated (Figures 4C and S3B). Visually, this corresponds to “fast lanes” that transport low-c-di-GMP cells towards the periphery during biofilm development (Video S2).

To further establish a causal relation between c-di-GMP phenotype and cell dynamics during biofilm growth, we engineered binary populations of c-di-GMP-based cell-types, in which cells switch states irreversibly via stochastic plasmid loss (Figure 4F). In the high-to-low-c-di-GMP scenario, WT cells harboring a plasmid that encodes a DGC (leading to high [c-di-GMP]) were grown in minimal medium plus exogenously supplemented spermidine *without antibiotics*; in these conditions, a cell that stochastically lost the plasmid would irreversibly transition to a low-c-di-GMP state.^39^ We found that the low-c-di-GMP cells that emerge in these initially high-c-di-GMP populations translocate to and dominate the periphery of the biofilm, leading to a sorted pattern similar to that in WT biofilms (Figure 4G, top; Video S4). In the low-to-high-c-di-GMP scenario, we used a high-c-di-GMP background strain (*vpvC*^W240R^)^40^ that harbors a plasmid encoding a PDE. Here, we found that stochastic plasmid loss gives rise to the sporadic emergence of high-c-di-GMP cells in an initially low-c-di-GMP population, and these high-c-di-GMP cells act as “roadblocks” that do not move outward (Figure 4G, bottom; Video S4). Together with the lineage tracing data, these results demonstrate how c-di-GMP level dictates the dynamics of individual cells, and consequently their positional fates, in developing biofilms.

### Agent-based modeling reproduces radially sorted c-di-GMP pattern via differential drag

The various patterns formed by c-di-GMP phenotypes in the matrix mutants (Figure 3) and the correlation between cell velocity and c-di-GMP level (Figure 4) led to our major hypothesis that the sorted pattern arises from *differential drag* experienced by low- and high-c-di-GMP cells during biofilm development: Since c-di-GMP signaling promotes matrix production, which gives rise to cell-cell and cell-surface adhesion, low- and high-c-di-GMP cells may experience different levels of adhesion and friction with their respective microenvironments as the biofilm grows and expands.^41,42^ We resorted to agent-based models (ABMs) to test our hypothesis.^28,43^

To develop a model of a growing population of cells with heterogeneous c-di-GMP levels, we first hypothesized that c-di-GMP switching can be described with a simple model in which each cell stochastically switches between low- and high-c-di-GMP states with mean lifetimes τ_L_ and τ_H_, respectively (Figure 5A).^44^ We first experimentally measured these lifetimes in quasi-2D colonies of Δ*vpsL* cells carrying both the riboswitch-based biosensor and a fast c-di-GMP sensor, cdGreen2 (Figure 5B).^45^ cdGreen2 increases its fluorescence intensity upon binding c-di-GMP, and therefore responds much more rapidly to changes in c-di-GMP level. In contrast, the fluorescence intensity of the riboswitch-based biosensor is governed by slower processes such as gene expression, protein degradation, and dilution due to cell growth. Therefore, we expected that cdGreen2 better reflects the instantaneous [c-di-GMP] in each cell, whereas the c-di-GMP-dependent physicochemical properties of biofilm-dwelling cells—such as adhesion to other cells and the surrounding matrix—are likely better reflected by the riboswitch-based biosensor, since changes in these properties similarly depend on slow biochemical reactions and physiological processes. The patterns of c-di-GMP phenotypes reflected by the two biosensors agree well in exponentially growing biofilms (Figures S5A–S5C). At the single-cell level, we found that *I*_cdGreen2_ exhibits dynamics akin to stochastic two-state switching, thus lending credence to our two-state model (Figures 5A–5C). By fitting a two-state hidden Markov model (HMM; Figure S5D)^46^ to the *I*_cdGreen2_ traces of 51 single-lineage trajectories between 2 and 10 h of growth (Figures 5C and S5E), we obtained mean state lifetime estimates of τ_L_ ≈ 2.14 h and τ_H_ ≈ 3.91 h (Figure 5D; Methods). We also observed more gradual changes in *I*_c-di-GMP_ after each switching event predicted by the HMM, consistent with our expectation regarding the two sensors (Figure S5F). The measured lifetimes are also in close agreement with population-based estimates from WT biofilms (Figures S5G–S5I; Methods).

**Fig. 5.**
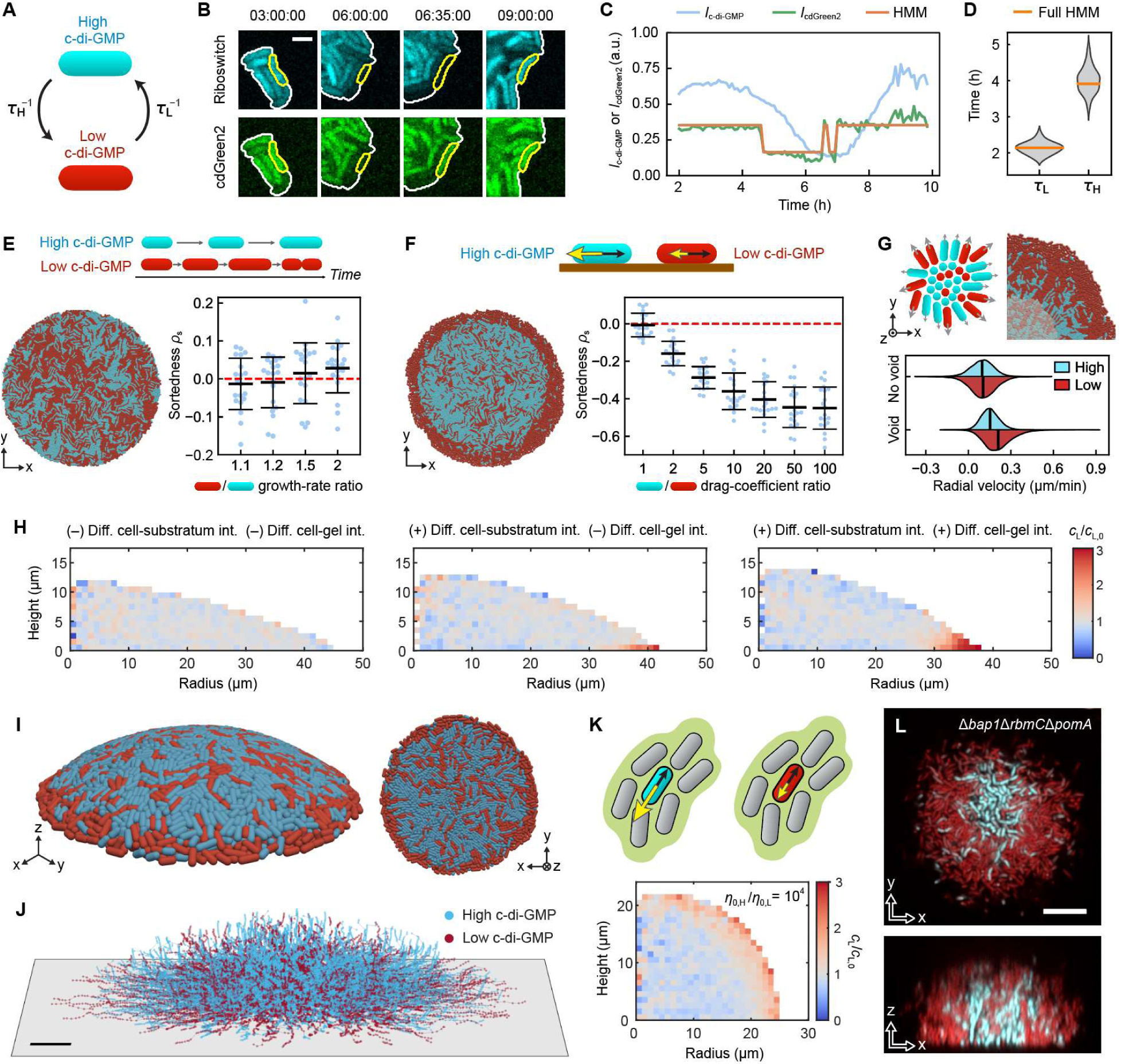
Agent-based modeling (ABM) of differential-drag-based sorting in biofilms. (**A**) Schematic of the model of stochastic switching of a cell between low- and high-c-di-GMP states, as a continuous-time Markov chain with mean state lifetimes τ_L_ and τ_H_ (thus, switching rates τ^-1^ and τ^-1^). (**B**) Snapshots of a single lineage in a Δ*vpsL* colony expressing both the riboswitch-based c-di-GMP biosensor (*top*, cyan) and a fast biosensor, cdGreen2 (*bottom*, green). (**C**) *I*c-di-GMP (cyan) and *I*cdGreen2 (green) in a single lineage from 2 to 10 h. Shown in orange are the state assignments, corresponding to low- and high-c-di-GMP states, predicted by a trained hidden Markov model (HMM). (**D**) Estimates of the mean state lifetimes τ_L_ and τ_H_ from an HMM fitted to 51 single-lineage trajectories, yielding τ_L_ ≈ 2.14 h and τ_H_ ≈ 3.91 h (orange); fitting the HMM to random subsamples of 40 trajectories each yielded a range of similar estimates (grey). (**E**) Growth-rate difference does not lead to radial sorting. (**F**) Radial sorting due to differential drag in the 2D ABM. In (E) and (F): *Top*: Schematic of a scenario in which low- and high-c-di-GMP phenotypes exhibit different growth rates due to metabolic cost (E) or differential drag due to matrix production (F). *Bottom left*: Example simulation outcomes of the spatial distribution of low- and high-c-di-GMP phenotypes. *Bottom right*: Sortedness, *ρs*, as a function of the ratio of growth rates (E) or of generalized drag coefficients (F; Document S1) between the two states. Both state lifetimes were set to 8 hours (see Figures S6A and S6B for data pertaining to other values). Error bars indicate mean ± s.d. (20 simulations). (**G**) *Top left:* Schematic of radial cell alignment due to the presence of a central growth void. *Top right:* An example simulation with a 20-fold difference in drag and a growth void (shaded region) whose radius was set to half of the biofilm radius (Document S1). *Bottom*: the radial velocities of low- and high-c-di-GMP cells in simulations with a growth void show a greater difference than in simulations with no void, due to the emergence of “fast lanes” (Document S1). (**H**) Distribution of c-di-GMP phenotypes, shown as the local relative fraction of low-c-di-GMP cells, *c*_L_/*c*_L,O_, in the presence or absence of differential interactions along the substratum or the biofilm-gel interface in 3D ABMs. Here, *c*_L_ is the local fraction of low-c-di-GMP cells and *c*_L,O_ ≡ τ^-1^/(τ^-1^ + τ^-1^) is the expected steady-state fraction of low-c-di-GMP cells across the biofilm. Data are shown as (*r*, *H*) projections averaged over 10 simulated biofilms (τ_L_ = 2 h and τ_H_ = 4 h). (**I**) Example simulation outcome from the 3D ABM, containing 8294 cells at 10 h, incorporating both differential cell-substratum and cell-gel friction, shown in a 3D perspective view (*left*) and from the bottom (*right*). (**J**) Single-lineage trajectories from the same simulation shown in (I), color-coded by the instantaneous c-di-GMP phenotype. 1000 randomly sampled trajectories between 0 and 10 h are shown. (**K**) Differential cell-cell adhesion, modeled as difference in ambient viscosity, leads to sorting in both *r* and *z* directions. *Top*: Schematic of differential viscous force (yellow arrow) experienced by a low- and high-c-di-GMP cell moving through the matrix and between neighboring cells at the same velocity (black arrow). *Bottom*: Distribution of c-di-GMP phenotypes, shown as the relative fraction of low-c-di-GMP cells (*c*_L_/*c*_L,O_), averaged over 10 simulations wherein the high-c-di-GMP cells experience a 10000-fold greater dynamic viscosity than the low-c-di-GMP cells. (**L**) Confocal images of the bottom layer (*top*) and vertical cross-section (*bottom*) of a Δ*bap1*Δ*rbmC*Δ*pomA* biofilm embedded in a soft hydrogel (0.225%), showing sorting in both *r* and *z* directions. The non-motile mutant (Δ*pomA*) was used to prevent cells from escaping into the soft gel. Scale bars: 2 μm in (B) and 10 μm in (J, L).

We then extended a previously described ABM of biofilm growth to incorporate switching between the low- and high-c-di-GMP states (Methods; Document S1).^27,28,43,47^ In this ABM, each cell is modeled as a spherocylinder that grows exponentially and divides upon reaching a critical length. The cells are also subject to mechanical forces with each other, the glass substratum, and the agarose gel, which is modeled as a coarse-grained network of spherical particles (Figure S6C). Moreover, the motion of each cell is counteracted by drag due to friction against the substratum and/or gel, as well as ambient viscosity within the bulk of the biofilm. We also assume that switching between the low- and high-c-di-GMP states proceeds with rates that are independent of spatial position or time. Using different versions of this ABM, we sought to identify putative phenotypic differences between the low- and high-c-di-GMP states that can give rise to the sorted pattern.

We first implemented a simplified set of 2D ABMs and considered two hypothetical sorting mechanisms: (1) high-c-di-GMP cells grow more slowly than low-c-di-GMP cells, due to the metabolic burden of matrix production; and (2) high-c-di-GMP cells experience greater drag than low-c-di-GMP cells, since the matrix facilitates adhesion to surfaces and other cells. First, we found that imposing differential growth rates fails to give rise to sorting, even when the growth rates differ by a factor of 2 (Figures 5E and S6A). This differs from observations in colonies grown on agar plates, in which strains with growth advantages dominate the expansion front;^48,49^ our ABMs show that differential growth does not give rise to sorting when the growth-rate difference arises from stochastic switching in a spatially uniform manner. In contrast, differential drag *does* give rise to sorting (Figures 5F and S6B). Moreover, the sorted pattern becomes more pronounced (ρ_s_ becomes more negative) as the ratio between the drag coefficients of the two states increases. Sorting was observed for all combinations of mean state lifetimes that we considered (Figure S6B), albeit to varying extents: Generally, increasing either state lifetime strengthened the pattern. Intuitively, the longer the cells can sustain their phenotype, the more time is available for the low-c-di-GMP cells to overtake the high-c-di-GMP cells towards the expansion front before they switch. We were also able to reproduce radial cell organization and the formation of fast lanes by introducing a core of non-growing cells, which mimic the verticalized cells in 3D biofilms that do not contribute to in-plane expansion of the bottom layer (Figure 5G).^27^ In all, these results support our hypothesis that differential drag is a viable physical mechanism for sorting.

While the simplicity and computational efficiency of our 2D ABMs enabled preliminary hypothesis testing of sorting mechanisms, they simplify the various types of physical interactions experienced by the cells and are inadequate for capturing new features that emerge in three dimensions. Therefore, we developed a full 3D ABM that incorporates the two-state model. We first tested 3D ABMs in which differential cell-surface adhesion between the low- and high-c-di-GMP cells generates differential friction at the interfaces with the glass substratum and the gel. These models revealed that imposing differential cell-substratum friction between the low- and high-c-di-GMP states gives rise to significant sorting in cell layers close to the base (Figure 5H, middle), compared to the null case (Figure 5H, left). Additionally imposing differential cell-gel friction strengthened the sorted pattern (Figures 5H, right, and 5I; see also Figure S6D). As with the 2D ABMs, we found that increasing state lifetimes in the 3D ABMs further strengthens the pattern (Figure S6E). Moreover, analysis of single-lineage trajectories in these simulations revealed that low-c-di-GMP cells tend to translate further outward in the radial direction than high-c-di-GMP cells (Figure 5J), consistent with the experimental results (Figures 4A and S3A). These findings further support differential friction as a mechanism for generating sorting in biofilms.

Finally, we sought to understand whether cell-cell adhesion can also give rise to sorting. Intuitively, cell-cell adhesion enables high-c-di-GMP cells to resist displacement by neighboring entities, an effect that can be phenomenologically modeled as an enhanced effective viscosity^50^ (Figure 5K, top). In contrast, since the low-c-di-GMP cells do not actively produce VPS, they cannot adhere to the surrounding matrix and/or cells,^42^ resulting in a decreased effective viscosity. Incorporating this differential viscosity into the 3D ABM, we found that the resulting biofilms exhibit radial sorting in 3D, i.e., a core of high-c-di-GMP cells surrounded by a shell of low-c-di-GMP cells (Figure 5K, bottom). Indeed, we can experimentally reproduce this 3D sorting by growing biofilms under a soft agarose gel (Figure 5L), which assume a more hemispherical geometry.

Together, the ABMs and ABM-guided experiments indicate that phenotypic sorting in biofilms arises from differential drag experienced by low- and high-c-di-GMP cells, generated by stochastic switching, as they are pushed outward by flows generated by the proliferation of other cells in the biofilm. This differential drag may arise from cell-substratum, cell-cell, cell-matrix, or cell-gel interactions, depending on the presence of various matrix components.

### Quorum sensing and nutrient limitation modulate the c-di-GMP pattern

As biofilms grow in size, diffusion becomes limited and chemical gradients are established along the radial direction^51^. One family of molecules that can exhibit such spatial gradients are autoinducers (AIs), which are secreted and detected by bacterial cells to infer local cell density, in a process known as quorum sensing (QS).^52^ In a densely packed biofilm, AI concentrations are expected to be high at the biofilm center and low at the periphery (Figure 6A). In *V. cholerae*, QS intersects with c-di-GMP signaling and controls biofilm formation: At high cell density, the master regulator HapR downregulates c-di-GMP by transcriptionally regulating VpsT and various DGCs and PDEs;^53^ HapR also directly downregulates matrix gene expression (Figure 6B).^54^ Therefore, we hypothesized that cells in the biofilm core may exhibit lowered c-di-GMP levels due to QS. Indeed, we consistently observed a dip in *I*_c-di-GMP_ close to the biofilm center (Figures 3A and 3B), which cannot be explained by the differential drag mechanism alone. To dissect the interplay between QS and c-di-GMP signaling, we generated a dual reporter strain harboring the c-di-GMP biosensor and a HapR-mNG fusion. Consistent with our hypothesis, HapR-mNG expression decays monotonically from the center to the edge, such that the dip in *I*_c-di-GMP_ coincides with regions with the highest *I*_HapR-mNG_ signal (Figures 6C, 6D, and S7). Following this logic, we hypothesized that deleting *hapR* would eliminate the dip in *I*_c-di-GMP_, resulting in monotonically decreasing *I*_c-di-GMP_ with radial position. This was indeed the case for the Δ*hapR* biofilms (Figures 6E and 6F). These results suggest that QS modulates the sorted pattern in a spatially dependent manner.

**Fig. 6.**
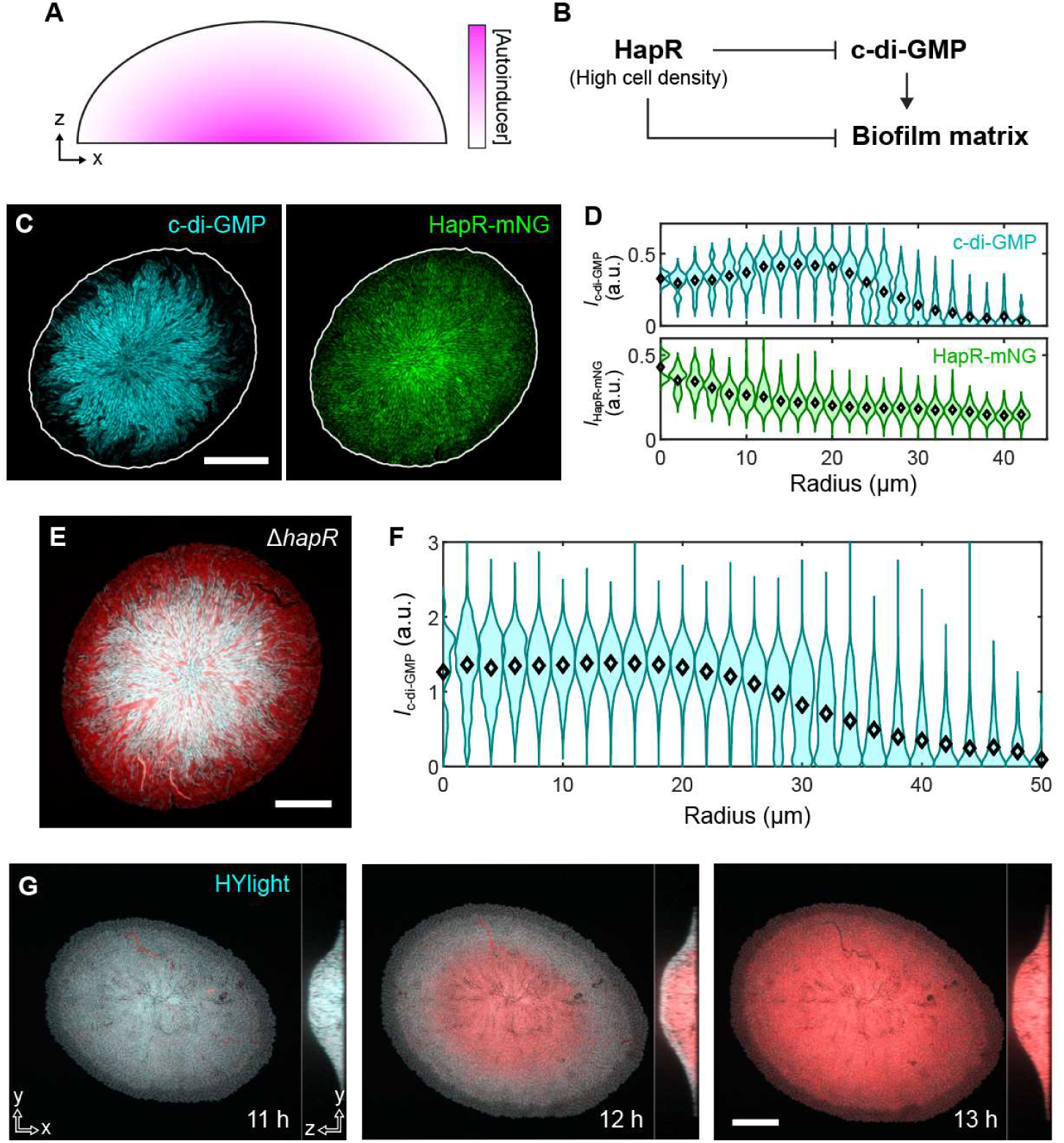
Quorum sensing (QS) and nutrient availability modulate the spatial patterning of c-di-GMP phenotypes. **(A)** Schematic of a biofilm’s vertical cross-section and the autoinducer (AI) concentration gradient due to secretion and diffusion. (**B**) Intersection of QS and c-di-GMP signaling in *V. cholerae*. (**C**) Confocal images of the bottom layer of a biofilm at 11 h expressing both the c-di-GMP biosensor (*left*) and HapR-mNG (*right*), showing a dip in c-di-GMP biosensor fluorescence and a corresponding increase in HapR-mNG fluorescence in the center. (**D**) Quantification of *I*c-di-GMP (*top*) and *I*HapR-mNG (*bottom*) in the bottom layer of the biofilm in (C). (**E**) Confocal image of the bottom layer of a biofilm at 11 h formed by a Δ*hapR* mutant expressing both a constitutive marker (red) and the c-di-GMP biosensor (cyan). (**F**) Quantification of *I*c-di-GMP in the bottom layer of Δ*hapR* biofilms. Data were obtained from 11 biofilms from 3 independent experiments, at timepoints at which each biofilm had a radius of ∼40 μm. Data in (D) and (F) are shown by violin plots and markers indicating the mean within each radial distance bin. (**G**) Confocal images of the bottom layer and vertical cross-section of a WT biofilm expressing HYlight, a reporter for glycolytic flux, showing decreased metabolism after 12 h. Scale bars: 20 μm.

Finally, we also found that the global decrease in [c-di-GMP] at later stages of biofilm development is facilitated by nutrient limitation. Using HYlight, a fluorescent reporter for intracellular abundance of fructose 1,6-bisphosphate, which serves as a proxy for glycolytic flux,^55^ we found that cell metabolism decreases sharply and globally after 12 h of growth (Figure 6G). This coincides with the global decrease in [c-di-GMP] (Figure 2B).

### c-di-GMP heterogeneity enhances adaptive fitness

Since low c-di-GMP is typically associated with the planktonic phenotype in *V. cholerae*, we hypothesized that the low-c-di-GMP cells that continuously emerge during biofilm growth are in the planktonic state. Indeed, when we removed the gel atop a confined biofilm, we found that the low-c-di-GMP cells at the periphery immediately departed, leaving behind the high-c-di-GMP cells at the center (along with some trapped low-c-di-GMP cells) (Figure 7A). This suggests that low-c-di-GMP cells in these biofilms are primed for motility. When we performed the same experiment without gel confinement—a more conventional growth condition—we instead observed mostly high-c-di-GMP cells close to the substratum (Figure 7B). We hypothesized that, without confinement, the low-c-di-GMP cells swim away from the substratum during the imaging intervals, creating a “survivor bias” where one only sees the surface-adhered subpopulation. Indeed, by growing and imaging WT biofilms in a microfluidic device and collecting cells in the liquid medium, we found that cells in the effluent are significantly lower in *I*_c-di-GMP_ compared to surface-adhered cells, suggesting a partitioning of high- and low-c-di-GMP cells on the surface and in the liquid medium, respectively (Figure 7C). To further support this idea, we found that removing cell motility restores the low-c-di-GMP subpopulation on the surface, as well as the spatial distribution of high- to low-c-di-GMP cells from center to periphery, in an unconfined geometry under non-flowing conditions (Figures 7D and 7E). These results suggest that c-di-GMP level is intrinsically heterogeneous in *V. cholerae* populations, independently of confinement, and low-c-di-GMP cells are sorted to the periphery of surface-attached biofilm clusters and continuously enter the bulk liquid medium as planktonic cells during biofilm growth.

**Fig. 7.**
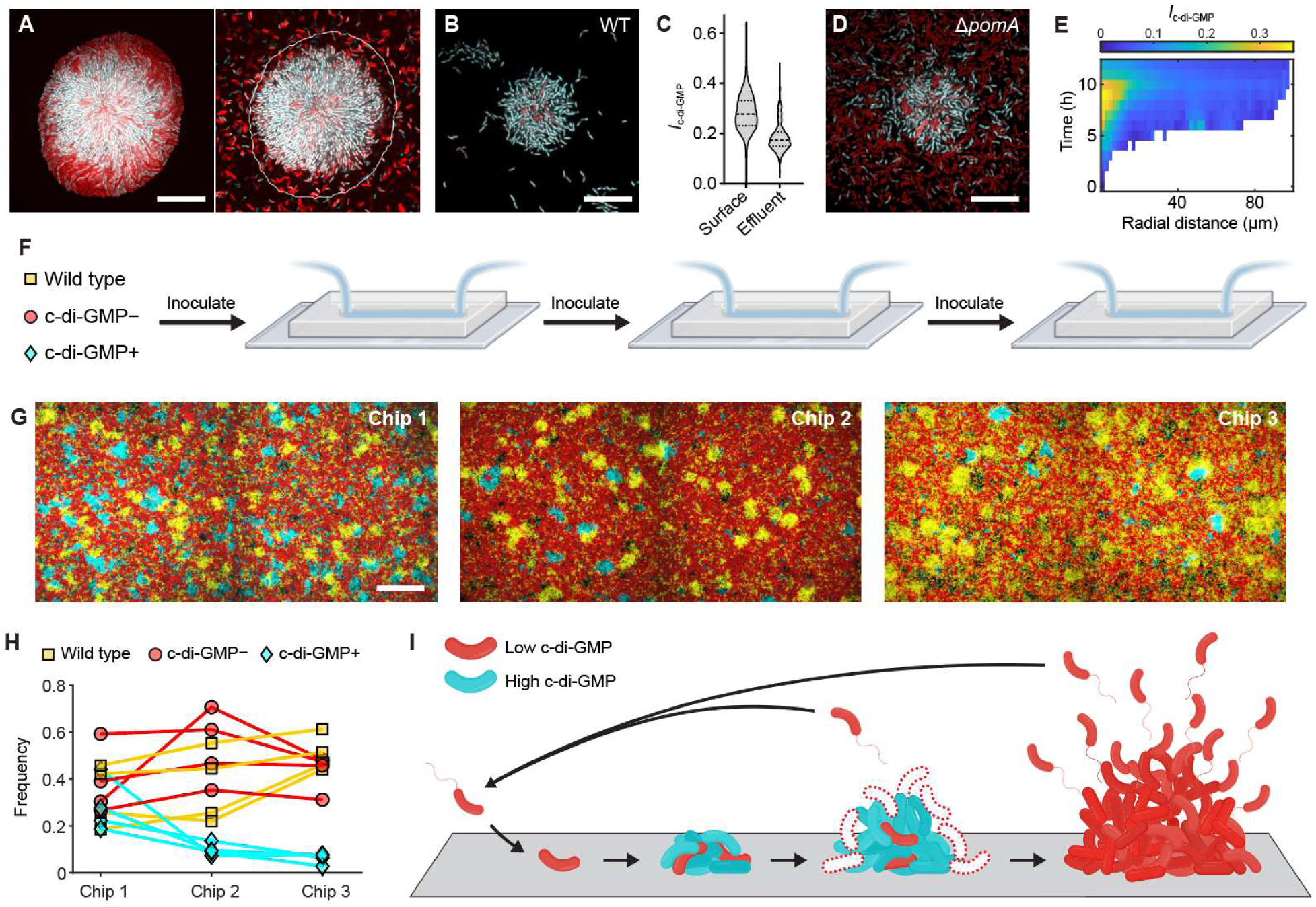
Heterogeneity in c-di-GMP signaling enhances adaptive fitness. (**A**) Confocal images of a WT biofilm under hydrogel confinement, before (*left*) and after (*right*) lifting the hydrogel, showing the departure of low-c-di-GMP cells at the periphery. (**B**) Confocal image of a WT biofilm grown without confinement. Low-c-di-GMP cells were not observed around the high-c-di-GMP cell cluster. (**C**) Comparison of *I*c-di-GMP between WT cells adhered to the glass substratum and those collected from the effluent of a microfluidic channel (data from 4 independent replicates). (**D** and **E**) Confocal image of a Δ*pomA* biofilm grown without confinement (D) and the quantification of *I*c-di-GMP in the bottom layer of the biofilm (E). (**F**) Competition between WT, c-di-GMP−, and c-di-GMP+ strains in a fluctuating environment emulated via consecutive microfluidic growth (6 h) and inoculation of each subsequent chip with the effluent from the previous chip. (**G**) Maximum intensity projections of confocal images of WT (yellow), c-di-GMP− (red), and c-di-GMP+ (cyan) cells in a representative channel in each of the three microfluidic chips. (**H**) Frequency of WT, c-di-GMP−, and c-di-GMP+ strains in the three microfluidic chips. Each set of connected data points represents an independent replicate. (**I**) Schematic of how heterogeneity in c-di-GMP signaling gives rise to the emergence of low-c-di-GMP cells during biofilm growth, which can leave the original biofilm cluster and colonize new niches before the final dispersal stage, thereby facilitating bet-hedging and improving colonization efficiency in uncertain and heterogeneous environments. Hollow cells with red dashed outlines indicate low-c-di-GMP cells that have been sorted to the biofilm periphery and have left the biofilm. Scale bars: 20 μm in (A, B, D) and 50 μm in (G). (F) and (I) were created using BioRender.com.

Phenotypic heterogeneity has long been associated with adaptive fitness in microbial communities in fluctuating environments through bet-hedging.^15^ To ascertain whether c-di-GMP heterogeneity confers a fitness advantage in *V. cholerae*, we performed a competition assay between WT *V. cholerae* and two other strains, c-di-GMP– and c-di-GMP+, which exhibit constitutively low and high c-di-GMP levels through a chromosomally integrated inducible PDE or DGC, respectively (Figure 7F; Methods). We inoculated the three strains with different fluorescent labels at equal density in a microfluidic chip, grew them for 6 h (still in the exponential phase), quantified the biomass of each strain with a confocal microscope, and used the effluent to inoculate the 2^nd^ chip. This process was repeated for the 2^nd^ and 3^rd^ chips. This experimental design simulates natural or host niches that harbor transient local hotspots for biofilm growth, and emerging planktonic cells can be carried by flow and colonize new niches. Results showed that the WT strain exhibits increased fitness with successive recolonizations, compared to the c-di-GMP– or c-di-GMP+ strains (Figures 7G and 7H). This advantage does not exist in static biofilm growth conditions, nor in pure liquid culture.^56^

## Discussion

In this study, by tracking the intracellular concentration of the key second messenger molecule c-di-GMP and downstream matrix production in growing biofilms at single-cell resolution, we have shown that biofilm-forming bacteria can exhibit strong phenotypic heterogeneity during biofilm development. In turn, the differential drag experienced by low- and high-c-di-GMP cells, which arises from differential surface friction and viscosity due to interactions with their micromechanical environments, facilitates phenotypic sorting as the biofilm expands. We have also shown how chemical signaling, including QS, can modulate this phenotypic patterning, in a manner reminiscent of signal-dependent cell-type differentiation in eukaryotic development.^57^ While phenotypic heterogeneity in dense biofilms has commonly been observed to arise from cells responding to local chemical environments predetermined by their positions,^16^ we show here the conceptually opposite scenario, in which heterogeneous gene expression stemming from a key intracellular phenotype (c-di-GMP level) determines each cell’s positional fate in the biofilm. This opposite scenario is more dominant in individual biofilm clusters, which are generally not large enough for the establishment of the significant oxygen or nutrient gradients that appear in mature, thick biofilms;^51^ biofilms of such size (tens of microns) may be more prevalent in nature and during infections for *V. cholerae*.^58–60^ Finally, we demonstrate that this heterogeneity continuously seeds a planktonic population, leading to a fitness advantage in fluctuating environments through a bet-hedging strategy.

It is instructive to compare these biofilms with eukaryotic developmental systems, in which different cell-types form patterns through mechanisms such as morphogen gradients,^61^ Turing-type reaction-diffusion dynamics,^62^ and oscillatory mechanisms such as the segmentation clock in vertebrate somitogenesis.^63^ The differential drag mechanism proposed in this study is perhaps most comparable to cell sorting via differential adhesion and/or interfacial tension between cell-types^20–23^. In general, sorting in these systems requires active mechanisms, such as motility, active neighbor exchange, or remodeling of the cortical cytoskeleton, which facilitate cellular reorganization in such a way that lowers the energy of the system. This principle also applies to sorting in bacterial aggregates mediated by differential active force generation via pili, as was observed in *Neisseria gonorrhoeae*.^64^ In the *V. cholerae* biofilms studied here, however, motility is not what generates sorting; rather, the cells within these biofilms are differentially advected by material flow arising from the proliferation of all cells within the community. This growth-dependent, nonequilibrium process has previously been shown to underlie reproducible cellular ordering and trajectories in constitutively biofilm-forming strains.^27,28,38^ Here, the same growth-induced flow leads to spatial sorting of phenotypes in biofilms that exhibit strong heterogeneity, adding to the catalogue of mechanisms for pattern formation in natural and synthetic multicellular systems.^20–23,65^ We also note that, whereas eukaryotic cells often differentiate into terminal cell-types, each biofilm-dwelling cell retains the capacity to reversibly transition between phenotypic states—another key difference in the developmental trajectories of these systems.

An immediate question stemming from our observations is the *origin* of heterogeneity in c-di-GMP signaling in *V. cholerae*. While second-messenger-based phenotypic heterogeneity in clonal microbial populations has been widely observed,^66–68^ the molecular origins of such heterogeneity have remained unclear in many contexts. In *Caulobacter crescentus* and *Pseudomonas aeruginosa*, asymmetric cell division has been shown to result in c-di-GMP heterogeneity,^69,70^ but such a mechanism has not yet been identified in *V. cholerae*. One appealing possibility is the involvement of motifs such as feedback and feedforward loops—which are often invoked as mechanisms for generating bistability, heterogeneity, and oscillations^14,71^—in the c-di-GMP regulatory network. For example, the c-di-GMP-dependent master biofilm regulators VpsR and VpsT can upregulate DGCs and PDEs, thereby generating positive and negative feedback on c-di-GMP.^72–74^ Indeed, a positive feedback loop has been shown to engender c-di-GMP heterogeneity in *P. aeruginosa*.^75^ Moreover, fluctuations in c-di-GMP signaling may arise from other mechanisms such as zero-order ultrasensitivity,^76^ which may emerge when enzymes with opposing functions (here, DGCs and PDEs) are saturated by their substrates (here, GTP and c-di-GMP). In this regime, small changes in DGC/PDE concentrations due to molecular fluctuations or environmental cues may cause large changes in c-di-GMP levels, which would manifest as phenotypic heterogeneity across the community. Given the broadly significant regulatory roles of c-di-GMP, important work remains in pinpointing the critical components within its regulatory network that give rise to this heterogeneity.

The differential drag mechanism is rooted in the distinct physicochemical interactions between high/low-c-di-GMP cells and the matrix. While extracellular matrix has often been described as a “public good” shared among biofilm-dwelling cells,^77^ recent studies in *V. cholerae* have demonstrated that only cells that actively produce VPS can adhere to each other and to the matrix;^42^ in other words, VPS acts as a recognition mechanism between matrix-producing cells. Our observations are consistent with this picture, in which low-c-di-GMP cells minimally interact with the matrix and neighboring cells, thus giving rise to differential drag between low- and high-c-di-GMP cells.

Our findings from the competition assay suggest that the low- and high-c-di-GMP phenotypes jointly confer a bet-hedging strategy that balances the benefits of colonizing local surfaces and exploring new territories, improving fitness in uncertain and heterogeneous environments. Importantly, our picture deviates from the classical scheme in which cells coordinately disperse due to nutrient limitation or other environmental cues:^2,78^ Even during exponential growth, planktonic subpopulations are continuously generated from the biofilm through fluctuations in c-di-GMP signaling (Figure 7I). This planktonic subpopulation is generally missed in common experimental settings due to the inability to visualize it in an open geometry. Mechanistically, this bet-hedging is possible because, unlike *collective* signaling processes such as QS, c-di-GMP is an intracellular signal that is more susceptible to large molecular fluctuations within *individual* cells. The differential drag mechanism ensures that the low-c-di-GMP cells are preferentially transported to the periphery and thus primed to leave. This is potentially related to another key observation in *V. cholerae* biofilms: Dispersal takes place at the biofilm periphery instead of the core,^79^ opposite to what is observed in *P. aeruginosa*.^78,80^ Finally, we note that recent studies have also observed the coexistence of planktonic and surface-attached subpopulations in *Vibrio* species grown in *in vitro* analogues of marine environments with nutrient hotspots.^81,82^ Our results highlight c-di-GMP fluctuations as a potential key molecular mechanism that may facilitate this population partitioning. More broadly, we predict that the partitioning of cells into biofilm and planktonic states within the metapopulation also depends on various environmental and physiological factors—including nutrient availability, metabolic mechanisms, and predation—all of which can modulate the optimal switching strategy and, in turn, shape the evolution of the underlying regulatory network.^81–83^

Finally, we highlight several new experimental techniques that we have introduced here for studying biofilms. For example, we have developed the first experimental and analysis platform for tracing single lineages and their gene expression in 3D biofilms. While intracellular puncta have been previously used to map the flow pattern and positional cell fates in biofilms via light-sheet microscopy,^38^ simultaneously tracking individual cells and their gene expression had remained a formidable task. By combining the same bright puncta with an *asynchronous* multichannel imaging protocol, we succeeded in tracking the gene expression of single cells within a biofilm and generating partial lineage information, which had previously not been possible. Future work inspired by this strategy should eventually lead to a full lineage tracing protocol for growing 3D biofilms. Another example is the tetracysteine epitope for *in situ* labeling of matrix proteins in living biofilms. While fluorescent antibodies have often been used to visualize distributions of matrix proteins, their large size (∼10 nm) renders penetration an ever-present issue in large biofilms. The low-molecular-weight and fluorogenic biarsenical dyes circumvent this issue, allowing us to visualize the distribution of RbmA within mature biofilms. Moreover, the TC-labeling strategy is orthogonal to, and can be combined with, antibody labeling. Future advancements in single-cell imaging and matrix labeling tools—potentially combined with spatial transcriptomics^11,13^—will unveil other forms of phenotypic diversity in other model species or even in naturally occurring multispecies biofilms.

## Acknowledgements

We thank Dr. Fitnat Yildiz for illuminating discussions and the original plasmid containing the riboswitch-based biosensor. We thank Dr. Urs Jenal for sharing the cdGreen2 plasmid. We thank Dr. Hesper Rego for sharing the HYlight reporter, and Lucas Soares and Dr. Ronald Breaker for help with the plate reader. We thank Dr. Andrew Bridges for providing the HapR-mNG construct. We thank Drs. Scott Holley, Corey O’Hern, and Carey D. Nadell for helpful discussions. Research reported in this publication was supported by the National Institute of General Medical Sciences of the National Institutes of Health under Award Number DP2GM146253 (J.Y.). This publication was made possible in part with the support of the Charles H. Revson Foundation 22-28 (J.N.). J.-S.B.T. is a Damon Runyon Fellow supported by the Damon Runyon Cancer Research Foundation (grant no. DRG-2446-21).

## Author Contributions

J.-S.B.T., J.N. and J.Y. conceptualized the project. J.-S.B.T. constructed the bacterial strains and performed experiments. J.-S.B.T., K.-M.N., C.M.W. and J.Y. analyzed the data. K.-M.N., C.L. and S.Z. developed the simulation model. All authors contributed to the writing of the manuscript.

## Supplemental Information

Table S1. *V. cholerae* strains used in this study.

Table S2. Oligonucleotides used in this study.

Document S1. Description of agent-based models.

**Video S1. Time-course confocal imaging of a growing *V. cholerae* biofilm expressing a c-di-GMP fluorescent biosensor**. The video shows both the bottom layer and the vertical midplane cross-section of the biofilm. Fluorescence from the constitutive marker and the c-di-GMP biosensor are pseudo-colored red and cyan, respectively.

**Video S2. Time-course asynchronous multichannel imaging of a *V. cholerae* biofilm.** Fluorescence from the constitutive marker, c-di-GMP biosensor, and fluorescent puncta are pseudo-colored red, cyan, and yellow, respectively. Yellow arrows indicate the velocity vectors of tracked puncta, each projected onto the corresponding cross-sectional plane. The second part of the video shows a subset of traced single-lineage trajectories during biofilm development, color-coded by *I*_c-di-GMP_. Timecode denotes hours:minutes:seconds. Scale bar: 10 µm.

**Video S3. Example single-lineage trajectories illustrating the anti-correlation between c-di-GMP level and radial velocity.** In the first trajectory, an increase in *I*_c-di-GMP_ is accompanied by a halt in its outward motion. In the second trajectory, a decrease in *I*_c-di-GMP_ is accompanied by an increase in radial velocity, *v*_r_. Timecode indicates hours:minutes:seconds.

**Video S4. Engineered binary populations reveal a causal relation between c-di-GMP level and cell dynamics.** Time-course confocal images of the bottom layers of two biofilms show individual cells undergoing irreversible, stochastic switching in intracellular c-di-GMP level due to plasmid loss. The left panel shows transitions from high to low c-di-GMP, and the right panel shows transitions from low to high, both resulting in corresponding changes in cell dynamics. Timecode indicates hours:minutes:seconds.

## Methods

### Bacterial strains and growth conditions

All *V. cholerae* strains used in this study are derivatives of the wild-type *Vibrio cholerae* O1 biovar El Tor strain C6706str2 and listed in Table S1. Mutations were genetically engineered using the natural transformation method (MuGENT) or through the suicide vector pKAS32.^84,85^

The riboswitch-based fluorescent c-di-GMP biosensor was designed based on a triple-tandem c-di-GMP riboswitch (*Bc3*, *Bc4*, and *Bc5* RNA) in *Bacillus thuringiensis* subsp. *chinensis* CT-43.^24^ The riboswitches assume transcriptional terminator conformations in the absence of c-di-GMP, but allow for downstream transcription when bound to c-di-GMP. We cloned the sequentially arranged *Bc3* and *Bc4* riboswitches from a c-di-GMP reporter plasmid and inserted it between a P*_tac_* promoter and a degradable mNG containing an ASV tag (AANDENYAASV) at the C-terminus.^86^ This construct was stitched together with a copy of P*_tac_*-mScarlet-I for cell labeling, with a synthetic transcriptional terminator L3S2P21 in between,^87^ and inserted into the chromosome at VC1807, a neutral locus in *V. cholerae* (Figure S1A).^88^ P*_tac_* functions as a constitutive promoter in *V. cholerae* C6706. Additional reporters were inserted into VC0501, another neutral locus in *V. cholerae*.^88^

Confined biofilm growth experiments were performed in M9 minimal media supplemented with 2 mM MgSO_4_, 100 μM CaCl_2_, 0.25% glucose, and 0.25% casamino acids (referred to as M9 media hereafter). Cells were first grown under shaken conditions overnight in LB broth (BD) at 30℃. The overnight culture was back-diluted 30× in M9 media and grown under shaken conditions at 30℃ until the optical density (OD_600_) reached 0.5–0.8 (about 3 h). The bacterial culture was diluted in M9 media without glucose or casamino acids to an OD_600_ of 0.001–0.003, and a 1 µL droplet of this diluted culture was deposited in the center of a glass-bottomed 96-well plate (MatTek). Concurrently, agarose polymer of a given concentration (1.5 wt% unless otherwise specified) was boiled in M9 media without glucose or casamino acids, and subsequently introduced into a mold made of glass slides and cover glass to solidify into a 170-μm-thick, approximately 4-mm-by-4-mm membrane. The membrane was then placed on top of the droplet of the diluted culture. A home-made silicone device that provides both aeration and moderate compression, dubbed “Snorkel,” was inserted into the well, pressing the agarose membrane and sandwiching the cells between the agarose membrane and the glass substratum (Figures S1C and S1D). Finally, 150 µL of M9 media was added to the well to supply nutrients. Cells were grown under static conditions at 27°C and imaged at various timepoints during growth using an adaptive imaging volume.^28^ For experiments in which the agarose atop a confined biofilm was lifted, the Snorkel was removed from the well and the agarose membrane spontaneously detached.

Unconfined biofilm growth experiments were conducted with the same growth conditions as in confined biofilm growth experiments, except without the confining agarose membrane. A Snorkel of a different size (see below) was used to ensure adequate aeration while maintaining an ∼100 μm gap between the Snorkel and the glass substratum.

### Snorkel manufacturing and sample geometry

Snorkels were manufactured with silicone to allow aeration and confinement of cells, as well as easy fitting to 96-well plates. Specifically, a 4-mm-by-4-mm and 127-μm-thick silicone membrane (McMaster-Carr) was placed at the bottom of a 96-well plate, and two 1/2” long and 1/8” diameter rod-shaped Neodymium magnets (McMaster-Carr) were used to sandwich the glass bottom and the silicone membrane and hold the membrane in place. Polydimethylsiloxane (PDMS, Sylgard 184, The Dow Chemical Company) silicone was filled into the well and cured at 55°C for 3 h. The magnet was then removed from the cured PDMS and a 3D-printed hollow cylinder was inserted into the space left behind, providing structural support and easy handling. Finally, extra PDMS was removed from the waist of the prototypical Snorkel with a scalpel to create space for a nutrient reservoir. To exert minimal compression on confined biofilms, two extra layers of 127-μm-thick silicone membrane were fused to the bottom of each Snorkel by curing PDMS, whereas Snorkels without these extra silicone membranes were used for unconfined biofilm growth. The as-manufactured Snorkels geometrically conform to the shape of the 96-well plate and provide confinement and aeration to cells through the silicone membrane (Figures S1C and S1D).

### Image acquisition and analysis

Imaging was performed using a Yokogawa CSU-W1 spinning-disk confocal scanning unit mounted on a Nikon Ti2-E microscope body, using the Nikon perfect focus system, and images were acquired by an sCMOS camera (Prime BSI, Teledyne) using Nikon Elements 5.20. For high-resolution, single-cell imaging, a 100× silicone oil immersion objective (Lambda S 100XC Sil, numerical aperture = 1.35) was used. Images of constitutive fluorescence, which were used for single-cell segmentation, were typically acquired at a *z*-step size of 0.130 μm, while the reporter fluorescence channels were acquired at a *z*-step size of 0.260 μm to reduce exposure. For time-course imaging, cells were incubated in a Tokai-Hit stage-top incubator set at 27°C and *z*-stack images were acquired at 1-h intervals. The height of time-course *z*-stack acquisition was automatically adjusted over time according to predetermined biofilm heights during growth.^28^ After acquisition, images were deconvolved using Huygens (version 20.04, SVI). The deconvolved image stacks were then segmented into individual cells using methods described in our earlier studies.^27,28,89^ Briefly, the images were first binarized layer by layer using a modified Bernsen’s local thresholding method, and the biofilms were then segmented into individual cells using an adaptive thresholding scheme. Each cell position and orientation were then determined from the center of mass and principal axis of the segmented voxels, respectively. We corrected for chromatic aberration between different channels using 0.1 μm multi-color fluorescent microspheres (TetraSpeck) and accounted for crosstalk using the crosstalk coefficients estimated by Huygens. The constitutive and reporter fluorescence intensities of the *i*-th segmented cell, FI_constitutive,i_ and FI_reporter,i_, were obtained as the average intensity of the segmented voxels of the cell in the constitutive and reporter channels, respectively. A calibrated relative fluorescence intensity, *I*_reporter,*i*_, was calculated as the relative fluorescence intensity (RFI) between FI_reporter,i_ and FI_constitutive,i_, calibrated by the *z*-variation in RFI (*f*(*z*)) based on biofilms of a control strain that constitutively expresses both reporters (see below) and additionally multiplied by the average FI_constitutive_ over all cells. Namely,

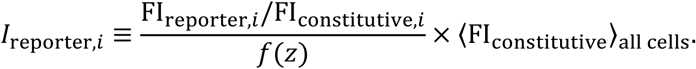

The RFI *z*-variation depends on the FP pair used and can be measured using biofilms of a control strain that constitutively expresses both the constitutive and reporter FP; the variation results from the different spectral properties and photostability of the two FPs when the confocal *z*-stack images are acquired from bottom to top. This *z*-variation exhibited reproducible patterns that were used to fit an exponential or polynomial function, depending on the reporter FP (Figure S1F), which was then used for calibration. Finally, we multiplied the *z-*calibrated RFI by the average FI_constitutive_ over all cells to obtain *I*_reporter,i_; this removes the contribution of 〈FI_constitutive_〉 at the population level, which, due to differences in the stability and expression dynamics of the constitutive and reporter FPs, can lead to spurious dynamics in *I*_reporter,i_. For example, FI_constitutive_ increases at later stages of biofilm growth due to growth-dependent accumulation of stable constitutive FPs, leading to small RFI values. To account for differing expression dynamics, in the time-course images shown in this study, the reporter channel is visualized using a fixed lookup table, while the constitutive channel is adjusted dynamically.

### Spatiotemporal visualization of *I*_reporter_

Spatiotemporal visualizations of *I*_reporter_ were generated by averaging single-cell *I*_reporter_ values within each spatiotemporal bin. For time vs. radius heatmaps of cells in the bottom layers of biofilms, cells within the bottom 1-μm layer of each biofilm at each timepoint were binned into 1–3 μm bins based on their distance from the biofilm’s origin, which was defined as the position of the founding cell. To agglomerate data from multiple biofilms, each bottom layer of a segmented biofilm at a given timepoint was transformed into a circular disk with radius equal to the average radius of all biofilms before binning and averaging. For height vs. radius heatmaps of agglomerated biofilm data at different timepoints, each segmented biofilm was first stretched in *z* to match the average height of all biofilms at each given timepoint. The biofilm was then divided into 1-μm *z* bins, and the layer of cells corresponding to each *z* bin was transformed into a circular disk with radius equal to the average layer radius at that *z* bin across all biofilms at that timepoint. This transformation was applied for all *z* bins and timepoints before averaging across biofilms, generating the final height vs. radius heatmaps.

### Sortedness quantification

The sortedness ρ_s_ of each biofilm was quantified using Spearman’s rank correlation coefficient between the radial distance from the biofilm’s origin and the *I*_reporter_ value of each cell in the bottom 1-μm layer. Quantification was performed on biofilms between 5 and 12 h of growth. The small number of cells prior to 5 h, and the low *I*_reporter_ signal after 12 h, made quantifications outside this time window less reliable.

### Staining of biofilm matrix

Immunostaining of the biofilm matrix protein RbmA was done using strains expressing C-terminal 3×FLAG-tagged RbmA under the same growth and experimental conditions as for time-course imaging experiments, except that the M9 medium contained Cy3-conjugated anti-FLAG antibodies (5 µg/mL; Sigma-Aldrich) and recombinant albumin (0.2 mg/mL; NEB); the latter blocks non-specific surface adsorption of fluorescent antibodies. Staining of RbmA by ReAsH-EDT_2_ was done using strains expressing RbmA tagged with an optimized tetracysteine (TC) motif (HRWCCPGCCKTF) at the C-terminus.^36^ M9 medium for RbmA-TC labeling additionally contained 10 μM of ReAsH-EDT_2_ labeling reagent (Invitrogen).

### Tracking cell dynamics and gene expression along single lineages by asynchronous multichannel imaging

Strains constitutively expressing mNG-μNS and containing the *Bc3*-*Bc4*-riboswitch-based c-di-GMP fluorescent biosensor were used to track cell position, velocity, and *I*_c-di-GMP_ along single lineages in biofilms. In these biofilm growth experiments, confocal images of each channel were obtained with different time intervals: at 3.75- or 5-min intervals, the bright intracellular mNG-μNS puncta were excited by a 488 nm laser and imaged at a *z*-step size of 0.260 μm, while, at 30-min intervals, both the mNG-μNS puncta and the constitutively expressed mScarlet-I FPs were imaged at a *z*-step size of 0.130 μm via a triggered acquisition setup that toggles between 488 nm and 561 nm laser excitation, and thus acquired both channels at each *z*-step, minimizing misalignment between puncta and cells due to biofilm growth. Additionally, SCFP3A fluorescence for *I*_c-di-GMP_ was imaged every 30 min at a *z*-step size of 0.260 μm. This asynchronous multichannel imaging setup ensures high time-resolution imaging of mNG-μNS puncta, required for single-lineage tracing, while minimizing light exposure leading to photobleaching and phototoxicity.

At each 30-min interval, the acquired 488 nm punctum images were thresholded using Nikon Elements to determine biofilm heights for live adaptive *z*-stack imaging, further minimizing laser exposure. The acquired images were first deconvolved by Huygens and registered using built-in functions in MATLAB to account for sample drift. Individual puncta were then identified as local maxima in each 3D confocal *z*-stack image, and subpixel localization was attained by fitting a paraboloid around each maximum. This process was repeated for images at all timepoints, and data for punctum coordinates at different timepoints were loaded into ImageJ and connected using the TrackMate particle-tracking software, using the LAP tracker with a penalty for linking based on punctum brightness, generating individual punctum trajectories.^90^ Because each mNG-μNS punctum was inherited by one daughter cell at division, the resulting punctum trajectory traces the position of a single lineage over time. The expression and assembly of new puncta in cells that did not inherit puncta after division allows the emergence of new single lineages. Instantaneous velocities along each trajectory were calculated using the central difference method and smoothed over a 22.5- or 30-min window. At the same time, the constitutive fluorescence images were segmented into individual cells and *I*_c-di-GMP_ was calculated using reporter fluorescence images as described above. To identify the cell hosting each punctum in a dense biofilm, each punctum’s coordinates were relaxed using a force field calculated as the gradient of the constitutive mScarlet-I image, until the punctum moved sufficiently towards the center of a segmented cell and an unambiguous assignment could be made. An alternative method that circumvents segmented-cell assignment, in which we calculated *I*_c-di-GMP_ within a spherical neighborhood of radius 0.325 μm around each relaxed punctum, yielded similar quantifications of *I*_c-di-GMP_ for each punctum. Thus, segmented single cells across different timepoints belonging to the same lineage were linked using the corresponding punctum trajectory, generating single-lineage trajectories that contain information on cell position, velocity, and *I*_c-di-GMP_ over time.

### Calculating correlation between *I*_c-di-GMP_ and *v*_*r*_ from single-lineage tracing data

To quantify the correlation between *I*_c-di-GMP_ and cell dynamics, we calculated its correlation with the relative deviation of *v*_r_, δ_v_ ≡ (*v*_r_ − *v*^-^_r_(*r*))/*v*^-^_r_(*r*), in the bottom layer. Here, *v*_r_ is the velocity of each tracked cell along the radial in-plane direction relative to the biofilm’s origin, and *v*^-^_r_(*r*) is the average radial velocity of all tracked cells at the same radius, *r*, as the cell. The latter was determined by binning the biofilm into 5-μm annuli from the origin and computing the mean *v*_r_ within each annulus. To quantify the correlation between *I*_c-di-GMP_ and *v*_r_ along the tangential direction (Figure S3B), *I*^-^_c-di-GMP_ and *v*^-^_r_ were calculated for each azimuthal angle as the average within the 30° angular sector centered at that angle.

### Engineering binary populations through stochastic plasmid loss

We engineered binary populations in which cells irreversibly switch c-di-GMP states upon losing a plasmid encoding either a DGC or a PDE. To generate a population in which cells first exhibit low c-di-GMP and stochastically switch to high c-di-GMP, we used a strain harboring the *vpvC*^W240R^ mutation that increases the activity of the DGC VpvC,^40^ resulting in constitutively high c-di-GMP. The strain was mated with a plasmid encoding an isopropyl ß-D-1-thiogalactopyranoside (IPTG)-inducible PDE, VC1086.^53^ We found, in this case, that cells retaining the plasmid exhibited constitutively low c-di-GMP. For the reverse scenario, a WT strain was mated with a plasmid carrying an IPTG-inducible DGC, *qrgB*.^53^ Both strains contain the chromosomal c-di-GMP biosensor, and both plasmids express SCFP3A for tracking plasmid retention. Confined biofilm growth experiments were performed as described above. Before biofilm growth, kanamycin was included in both LB and M9 media at 500–1000 μg/mL to provide selective pressure for plasmids, but no kanamycin was included during biofilm growth; instead, 1 mM IPTG was supplemented in the M9 medium to induce DGC or PDE expression. In the high-to-low-switching scenario, M9 medium contained 0.5% glucose, no casamino acids, and 1 mM spermidine to maintain constitutively low c-di-GMP in WT cells that have lost the plasmid.^39^

### *I*_c-di-GMP_ quantification of surface-adherent and effluent cells

To quantify *I*_c-di-GMP_ in surface-adherent cells and cells in the bulk liquid medium, four biological replicates of a WT strain with the riboswitch-based c-di-GMP biosensor were grown overnight at 30℃, back-diluted 30× and regrown in the M9 medium at 30℃ for 2 h, then back-diluted to OD_600_ = 0.03 before being introduced into the channels of a microfluidic chip (channel dimensions: length 1 cm, width 400 µm, height 60 µm) for 1 h to attach. After inoculation, sterile inlet and outlet polytetrafluoroethylene tubing (Cole-Parmer) were connected to the microfluidic chamber and M9 medium was flown through each channel at a flow rate of 0.1 µL/min controlled by a syringe pump (KD Scientific). After growing for 8 h at room temperature (∼25℃), z-stack confocal images at single-cell resolution of cells growing on the surfaces of the microfluidic channels of all four replicates were acquired. Effluents were collected after removing the outlet tubing, and cells in the effluent were sandwiched between an agarose pad and cover glass for confocal imaging. The confocal images were then used for *I*_c-di-GMP_ quantification.

### Quantifying *I*_c-di-GMP_ dependence on intracellular [c-di-GMP]

Intracellular [c-di-GMP] quantification was performed using a c-di-GMP Assay Kit (Lucerna, USA), based on the c-di-GMP-binding-dependent fluorescence from a c-di-GMP riboswitch and Spinach aptamer (Figure S1B).^91^ We used c-di-GMP biosensor strains carrying chromosomal arabinose-inducible constructs of a DGC or PDE and varied the arabinose concentration in the medium to vary the c-di-GMP level. The non-biofilm-forming Δ*vpsL* strain background was used to prevent clumping of cells and ensure accurate conversion between OD_600_ and colony-forming units (CFU). From overnight LB cultures, we back-diluted cultures of each strain to OD_600_ = 0.01 in M9 medium supplemented with different amounts of arabinose. The DGC/PDE constructs and arabinose concentrations used (in descending order of measured intracellular [c-di-GMP]) were: P*_BAD_*-*vpvC*^W240R^ arabinose 1%, P*_BAD_*-*vpvC*^W240R^ arabinose 0.25%, WT arabinose 0%, P*_BAD_*-VC1086 arabinose 0.25%, and P*_BAD_*-VC1086 arabinose 1%. Each strain was grown as two 15 mL cultures in two 50 mL centrifuge tubes with vigorous shaking for 6 h at 25℃ until OD_600_ reached ∼0.5. The cells were then harvested from the 30 mL cultures by centrifugation and resuspended in 560 μL cell lysis buffer, which included 100 μL BugBuster Protein Extraction Reagent (Novagen), 5 μL of 10 mg/mL lysozyme (Sigma-Aldrich), and 895 μL water for every mL of buffer. At the same time, a 300 μL culture of each strain was fixed with 4% paraformaldehyde (Electron Microscopy Sciences) for subsequent *I*_c-di-GMP_ quantification. After a 30-min incubation at room temperature, the lysates were incubated at 98℃ for 10 min to denature the FPs to reduce background fluorescence. Three replicates of 50 μL heat-denatured lysates, mixed with 150 μL of c-di-GMP assay master mix, along with the same mixture without the fluorophore or c-di-GMP biosensor in the master mix to control for residual FP signal and autofluorescence, were incubated at 25℃ overnight and quantified for fluorescence intensity in a 96-well clear-bottom plate using a plate reader (BioTek Synergy Neo2). The fluorescence intensity of the lysates, minus the background intensity from the controls, were used to calculate sample [c-di-GMP] based on a standard [c-di-GMP] vs. fluorescence curve. The sample [c-di-GMP] was converted to intracellular [c-di-GMP] by estimating the total intracellular volume of each culture using the following parameters: cell volume: 0.5 μm^3^, CFU per OD_600_ per mL: 1.37×10^9^. The measured *I*_c-di-GMP_ from the fixed cells and [c-di-GMP] from the quantification assay were combined to estimate the dependence of *I*_c-di-GMP_ on intracellular [c-di-GMP].

### Estimating lifetimes for low- and high-c-di-GMP phenotypes from single-lineage trajectories

To estimate lifetimes of low- and high-c-di-GMP phenotypes in *V. cholerae* cells, we grew Δ*vpsL* cells expressing both the riboswitch-based fluorescent c-di-GMP biosensor (*I*_c-di-GMP_) and a fast fluorescent c-di-GMP biosensor, cdGreen2 (*I*_cdGreen2_), into quasi-2D colonies in the confined geometry as described above. The colonies were imaged in the bottom layer at 5-min intervals, and the time-course images were subsequently deconvolved and registered to correct for sample drift. The processed 2D time-course images were then segmented using Omnipose, a general image segmentation tool useful for 2D images, using the constitutive fluorescence channel.^92^ We then manually traced 51 single lineages in these growing colonies using TrackMate in ImageJ.^90^ The constitutive and reporter fluorescence intensities of each cell at each timepoint were obtained based on the segmentation, from which we calculated *I*_c-di-GMP_ and *I*_cdGreen2_. Here, in calculating *I*_c-di-GMP_ and *I*_cdGreen2_, we ignored the *z*-variation correction *f*(*z*) and multiplication by the average FI_constitutive_ of all cells, since the Δ*vpsL* colonies were quasi-2D; moreover, we focused on a time window between 2 and 10 h in exponential phase, since the lifetimes are further affected by nutrient limitation and quorum sensing at later stages of biofilm growth.

To develop a model of phenotypic switching from these data, we trained a two-state hidden Markov model (HMM, Figure S5D).^46^ The model describes each cell as reversibly transitioning between two hidden states—corresponding to the low- and high-c-di-GMP phenotypes—according to a discrete-time Markov chain, with each transition occurring every 5 min, the time interval between images. We assume that, at each timepoint, the cell emits an observable fluorescence signal (namely *I*_cdGreen2_) according to a Gaussian emission distribution that depends on the hidden state (Figure S5D). To connect this to the two-state model (Figure 5A), we recall from the theory of continuous-time Markov chains that the infinitesimal transition rates, *k*_HL_ = τ^-1^ from high- to low-c-di-GMP and *k*_LH_ = τ^-1^ for *vice versa*, in the two-state model are given by, e.g.,^44^

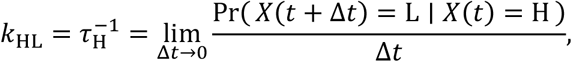

where Pr(⋅) denotes probability and *X*(*t*) is the state at time *t*. Therefore, assuming that Δ*t* = 5 min (which is the time interval between images) is sufficiently small, we can use the transition probabilities *P*_L,H_ and *P*_H,L_ in the HMM to approximate these rates (and the corresponding mean state lifetimes), as

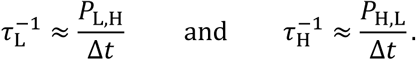

We implemented and trained this HMM in Python with the pomegranate package (version 1.0.0),^93^ using non-redundant segments from the above 51 single-lineage *I*_cdGreen2_ traces. Training the HMM on the entire dataset yielded mean lifetime estimates of τ_L_ = 2.14 h and τ_H_ = 3.91 h for the low- and high-c-di-GMP phenotypes, respectively; uncertainties for these estimates were quantified by re-training the HMM on random subsamples of 40 traces each (Figure 5D). The emission distributions of both hidden states were found to be highly peaked (low-c-di-GMP: mean ± s.d. = 0.166 ± 0.0390 a.u.; high-c-di-GMP: mean ± s.d. = 0.358 ± 0.0421 a.u.).

### Estimating lifetimes for low- and high-c-di-GMP phenotypes from population dynamics

We also followed an alternative approach to estimating the lifetimes of the low- and high-c-di-GMP phenotypes, by fitting Gaussian mixture models to *I*_c-di-GMP_ distributions obtained from 11 WT biofilms (Figures S5G–S5I). Upon visually inspecting the biofilm growth curves (Figure S5G), we chose a time window of 5 to 11 h optimized for this quantification, prior to which biofilms had very few cells (< 50), and after which most biofilms exhibited a slowdown in growth. We then fit the *I*_c-di-GMP_ distribution of each biofilm at each timepoint from 5 to 11 h to a two-component Gaussian mixture model,^46^ using the Python scikit-learn package (version 1.4.1; Figure S5H). This yielded estimates for the weights of the two Gaussian components, *w*_L_(*t*) and *w*_H_(*t*), over time in each biofilm. Moreover, we used the Bayesian information criterion to confirm that each of these *I*_c-di-GMP_ distributions was better described by a two-component Gaussian mixture than a pure Gaussian.

Let *P*_L_(*t*) (*P*_H_(*t*)) be the probability that the cell exhibits the low-(high-) c-di-GMP phenotype at time *t*, given that the cell’s phenotype evolves according to the two-state model, i.e., a two-state continuous-time Markov chain with transition rates *k*_LH_ = τ^-1^ and *k*_HL_ = τ^-1^ (Figure 5A). Then the time-evolution of these probabilities is given by the master equation,^94^

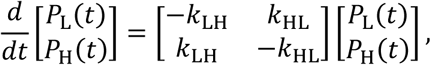

which admits the time-dependent solution,

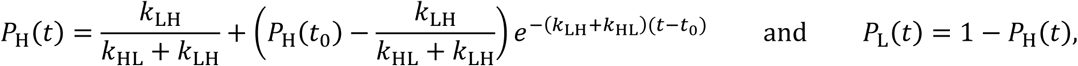

where *t*_O_ is an initial timepoint a_t_ wh_i_ch *P*_H_(*t*_O_) is known. Visually inspecting the *I*_c-di-GMP_ distributions, we found that almost all biofilms exhibit a uniformly high c-di-GMP _le_vel at 3 h, _a_nd _s_o we set *t*_O_ = 3 h and *P*_H_(*t*_O_) = 1. Finally, we fit the above solution for *P*_H_(*t*) to the avera_g_e weights of the high-c-di-GMP components over all 11 biofilms, {⟨*w*_H_(*t*)⟩: *t* ∈ [5 h, 11 h_]}_ (Figure S5I). This yielded estimates of *k*_LH_ ≈ 0.282 h^-1^ and _*k*HL_ ≈ 0.170 h^-1^, which corresponds to mean lifetimes of τ_L_ ≈ 3.55 h and τ_H_ ≈ 5.88 h.

### Agent-based simulations

The ABMs were built on those developed in previous work^27,28,43,47^ and updated to include cell-gel friction and the two-state model. For details, see Document S1.

### Competition assay using microfluidic channels

Competition experiments between WT, c-di-GMP+, and c-di-GMP– strains were performed using *V. cholerae* strains expressing (1) different FPs under a constitutive P*_tac_* promoter and (2) a DGC or PDE from chromosomal constructs under arabinose induction: P*_BAD_*-*qrgB* for c-di-GMP+, P*_BAD_*-VC1086 for c-di-GMP–, and P*_BAD_*-VC1086* (which encodes an inactive PDE)^53^ for WT to ensure that differences in metabolic burden due to enzyme production are minimized. Each strain was streaked onto an LB agar plate and grown overnight at 37℃. Single colonies of four biological replicates for each strain were picked and grown in LB shaken culture overnight at 30℃. The LB culture was then back-diluted 30× in M9 media and grown under shaken conditions until OD_600_ reached 0.2–0.3 after 2 h. The inoculant of the three competing strains was then mixed at equal proportions at OD_600_ = 0.05 and then introduced into the channels of the 1^st^ microfluidic chip (channel dimensions: length 1 cm, width 400 µm, height 60 µm) through the outlet. The cells were allowed to attach for 1 h, after which sterile inlet and outlet polytetrafluoroethylene tubing (Cole-Parmer) were connected to the microfluidic chamber and M9 medium supplemented with 0.25% arabinose was flown through the channels at a flow rate of 0.6 µL/min controlled by a syringe pump. The cells under flow were grown for 6 h at room temperature (∼25℃), before confocal images were taken with a 60× water immersion objective (CFI Plan Apo 60XC, numerical aperture = 1.20) at a *z*-step size of 1 μm for biomass quantification. Immediately after imaging, the outlet tubing was carefully removed, and effluents from the 1^st^ chip were collected and introduced into the 2^nd^ chip for inoculation. The same inoculation, growth, and confocal imaging procedures were repeated for the 2^nd^ and 3^rd^ chips. The biomass and frequency of each strain in each chip were quantified by binarizing the confocal images using a local thresholding method.^77^

## QUANTIFICATION AND STATISTICAL ANALYSIS

Errors correspond to standard deviations from measurements taken from distinct samples. Statistical analysis was performed using MATLAB (MathWorks) and Prism (GraphPad Software Inc.). Details of statistical analysis are included in figure captions.

