## Supplementary material for "Single-cell imaging reveals spontaneous phenotypic sorting and bet-hedging in developing biofilms": SI figures and Tables

### Supplementary figures

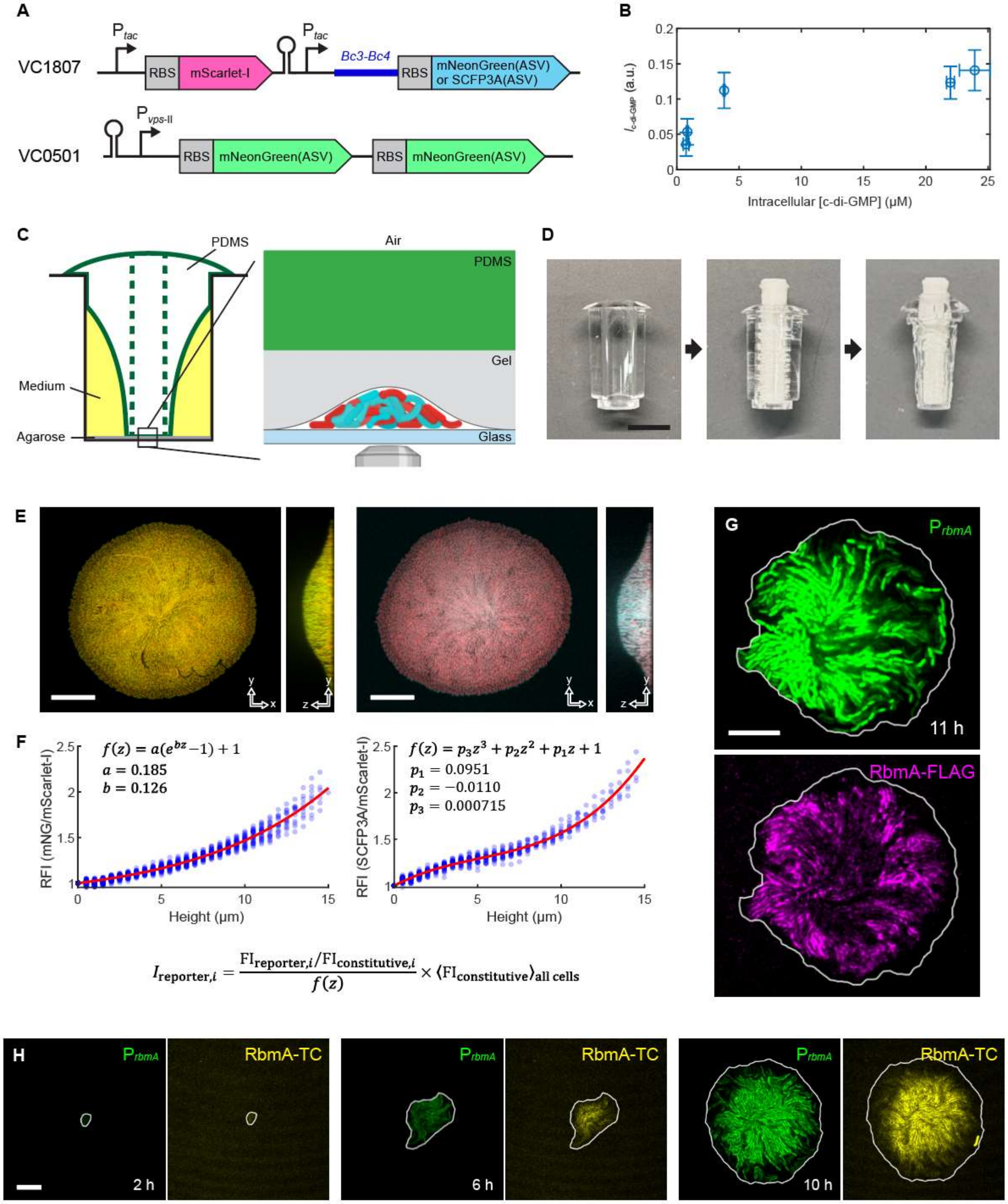

**Fig. S1. Strain construction, experimental systems, reporter characterization employed in this study, and biofilm matrix staining, related to Figures 1 and 2.**

(**A**) Construction of the riboswitch-based c-di-GMP biosensor and additional transcriptional reporters. Insertions were made at two neutral chromosomal loci, VC1807 and VC0501. In dual reporter constructs, SCFP3A(ASV) was used for the c-di-GMP biosensor instead of mNeonGreen(ASV).

(**B**) Quantification of the response of the *Bc3-Bc4* riboswitch at different intracellular [c-di-GMP]. $I_{c\text{-}di\text{-}GMP}$ was measured from *V. cholerae* cells in liquid cultures with different intracellular [c-di-GMP] via titrated induction of a

DGC or PDE. The bulk c-di-GMP level was measured using a c-di-GMP quantification assay (*Methods*). The response data provides the dynamic range of the c-di-GMP biosensor based on the *Bc3-Bc4* riboswitch, though $I_{\text{c-di-GMP}}$ values in liquid culture are generally lower than those in surface-attached biofilms. Error bars indicate mean ± s.d.

(**C**) Schematic of the PDMS "Snorkel" inserted into a well in a 96-well glass-bottom plate for aeration and confinement (*left*), and a close-up cross-sectional view of a biofilm under confinement (*right*). The drawings are not to scale. The thickness of the agarose gel and PDMS membrane in the experiments were 170 μm and 381 μm, respectively.

(**D**) Images of Snorkels and their manufacturing process. Scale bar: 5 mm.

(**E** and **F**) Calibration of single-cell $I_{\text{reporter}}$ using control strains expressing mScarlet-I and mNeonGreen(ASV) (E, *left*) or mScarlet-I and SCFP3A(ASV) (E, *right*) under the constitutive $P_{tac}$ promoter. The relative fluorescence intensity (RFI) in these control strains varies with height, and can be fitted by an exponential or cubic polynomial function, $f(z)$. These fits are shown as insets in (F). $I_{\text{reporter},i}$ was then calculated as the ratio between the reporter and constitutive fluorescence intensities, divided by $f(z)$ and multiplied by the average constitutive fluorescence of all cells. Data for fitting $f(z)$ for mNeonGreen(ASV) and SCFP3A(ASV) were collected from 20 and 8 biofilms, respectively. Scale bars: 20 μm.

(**G**) $P_{rbmA}$ activity and immunostaining of RbmA-3×FLAG in the extracellular space in a WT *V. cholerae* biofilm after 11 h of growth, showing ineffective penetration of fluorescently labeled antibodies in mature biofilms even when they are present throughout biofilm growth. As a consequence, the $P_{rbmA}$ activity and RbmA staining patterns are different. Scale bar: 10 μm.

(**H**) $P_{rbmA}$ activity and TC-tagged RbmA stained with a fluorogenic biarsenical dye, ReAsH-$EDT_2$, in a growing WT *V. cholerae* biofilm. RbmA-TC staining depends on covalent binding between RbmA-TC and ReAsH-$EDT_2$, which activates ReAsH fluorescence. A time delay was observed in TC labeling compared to FLAG-tag labeling. Scale bar: 10 μm.

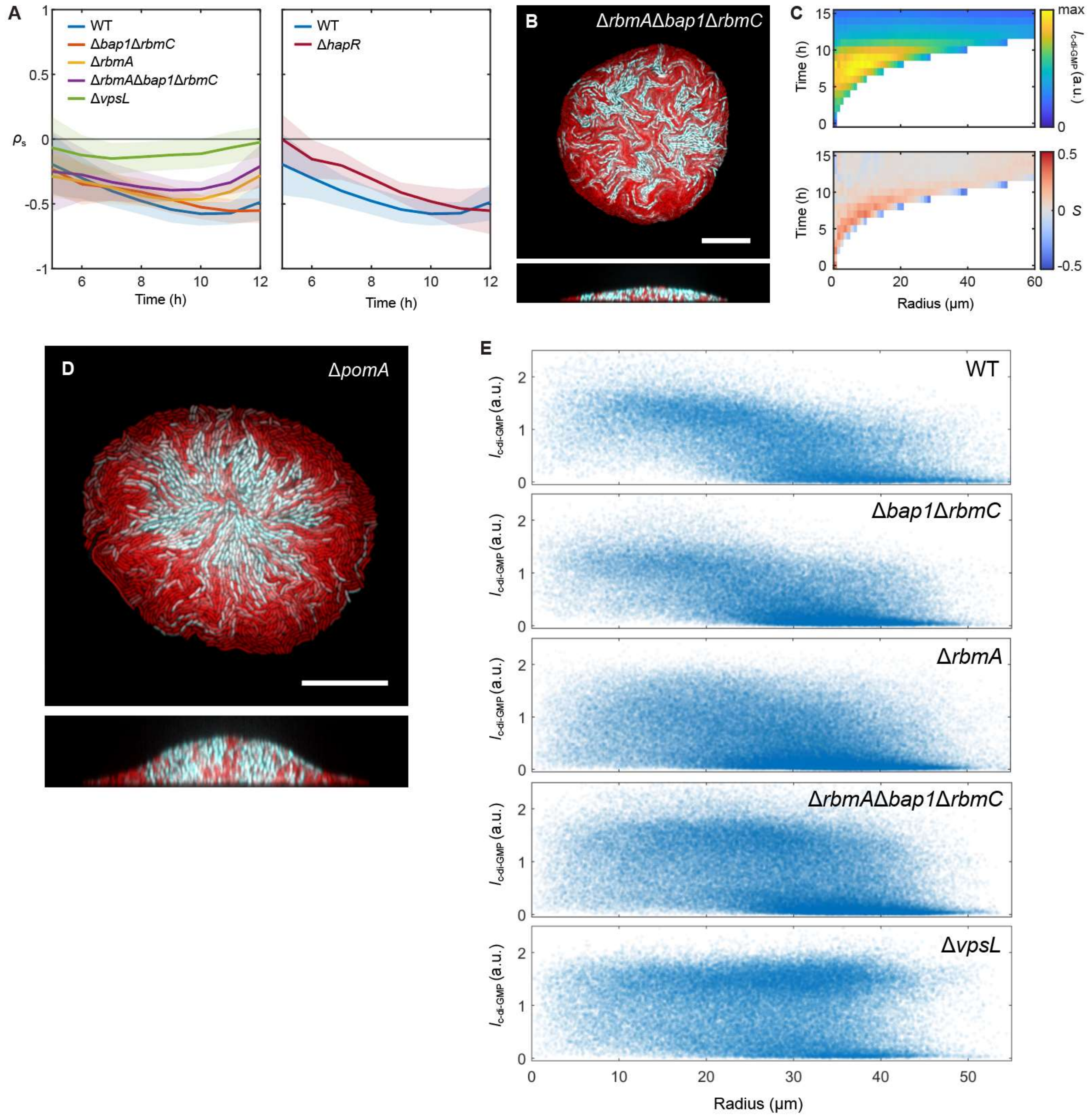

**Fig. S2. Sorting and radial alignment in WT and mutant *V. cholerae* biofilms, related to Figure 3.**

(**A**) $\rho_s$ vs. time for biofilms formed by WT *V. cholerae* and the matrix mutant strains Δ*bap1*Δ*rbmC*, Δ*rbmA*, Δ*rbmA*Δ*bap1*Δ*rbmC* and Δ*vpsL* (*left*), and biofilms formed by WT and the QS mutant strain Δ*hapR* (*right*). Solid lines and shading indicate mean ± s.d. Data for each strain were obtained from 11–25 biofilms from 3 independent experiments.

(**B**) Bottom layer (*top*) and vertical cross-section (*bottom*) of a *V. cholerae* biofilm formed by the matrix mutant strain Δ*rbmA*Δ*bap1*Δ*rbmC*, in which all three major matrix proteins are deleted but VPS is still properly synthesized. Scale bar: 20 μm.

(**C**) Time vs. radius heatmaps of locally averaged $I_{\text{c-di-GMP}}$ (*top*) and radial alignment, $S$ (*bottom*), of cells in the bottom layer of Δ*rbmA*Δ*bap1*Δ*rbmC* biofilms. Data were obtained from 20 biofilms from 3 independent experiments.

(**D**) Confocal images of a radially sorted biofilm formed by the non-motile Δ*pomA* mutant strain, showing that c-di-GMP-based sorting in confined biofilms does not depend on cell motility. Scale bar: 20 μm.

(**E**) Scatterplots of $I_{\text{c-di-GMP}}$ vs. radius of cells in the bottom layers of WT, Δ*bap1*Δ*rbmC*, Δ*rbmA*, Δ*rbmA*Δ*bap1*Δ*rbmC*, and Δ*vpsL* biofilms from 11–25 biofilms from 3 independent experiments for each strain. Timepoints were chosen so that the bottom layer contains ~ 5000 cells (radius of each biofilm was ~ 40 μm).

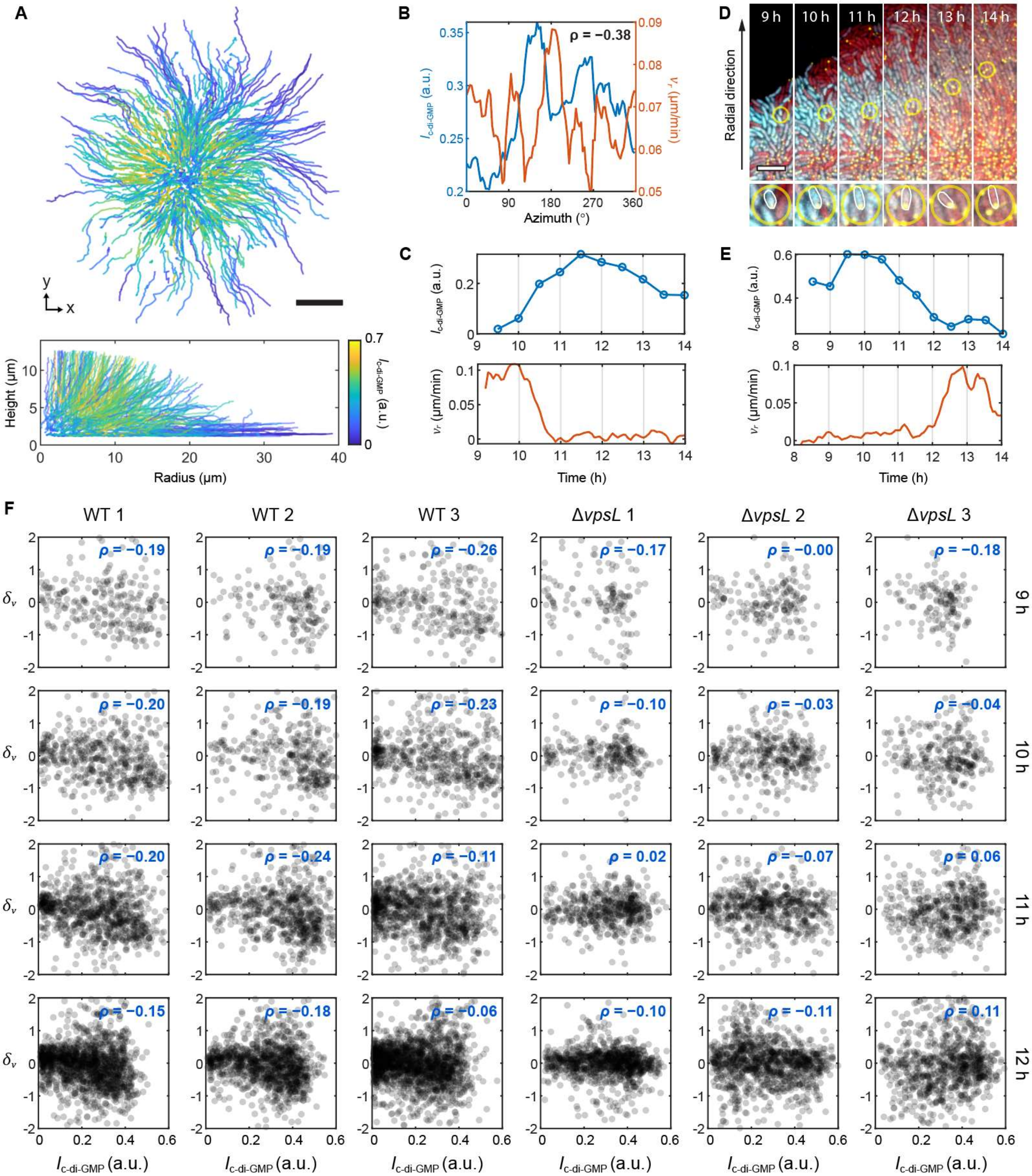

**Fig. S3. Single-lineage tracing reveals correlation between [c-di-GMP] and cell dynamics, related to Figure 4.**
(**A**) Single-lineage trajectories color-coded by $I_{\text{c-di-GMP}}$, shown as $(x, y)$ projection (*top*) and $(r, z)$ projection (*bottom*), within 12 h of biofilm growth. The 1000 and 500 trajectories that were tracked the longest in time are shown in the $(x, y)$ and $(r, z)$ projections, respectively. Scale bar: 10 μm.

(**B**) Radially averaged $I_{\text{c-di-GMP}}$ and radial in-plane velocity $v_r$ in the bottom layer of a biofilm after 11 h of growth, as functions of the azimuthal angle. These quantities exhibit a negative correlation, as quantified by Spearman's rank correlation coefficient. The radial averages at each azimuthal angle were taken within the 30° sector centered at that angle.

(**C**) Quantification of $I_{\text{c-di-GMP}}$ and $v_r$ along the trajectory shown in Figure 4D.

(**D** and **E**) Snapshots of an example single-lineage trajectory (highlighted by yellow circle) with decreasing $I_{\text{c-di-GMP}}$ and a corresponding change in cell position and velocity (D), as well as quantification of $I_{\text{c-di-GMP}}$ and $v_r$ along the trajectory (E). Scale bar: 5 µm.

(**F**) $\delta_v$ (relative deviation in $v_r$) versus $I_{\text{c-di-GMP}}$ of cells in the bottom layer of 3 WT biofilms and 3 Δ*vpsL* colonies, at different timepoints during biofilm growth. The Spearman's rank correlation coefficient, $\rho$, between $\delta_v$ and $I_{\text{c-di-GMP}}$ is given for each plot. WT biofilms consistently exhibited lower $\rho$ than Δ*vpsL* colonies, leading to the sorted c-di-GMP pattern.

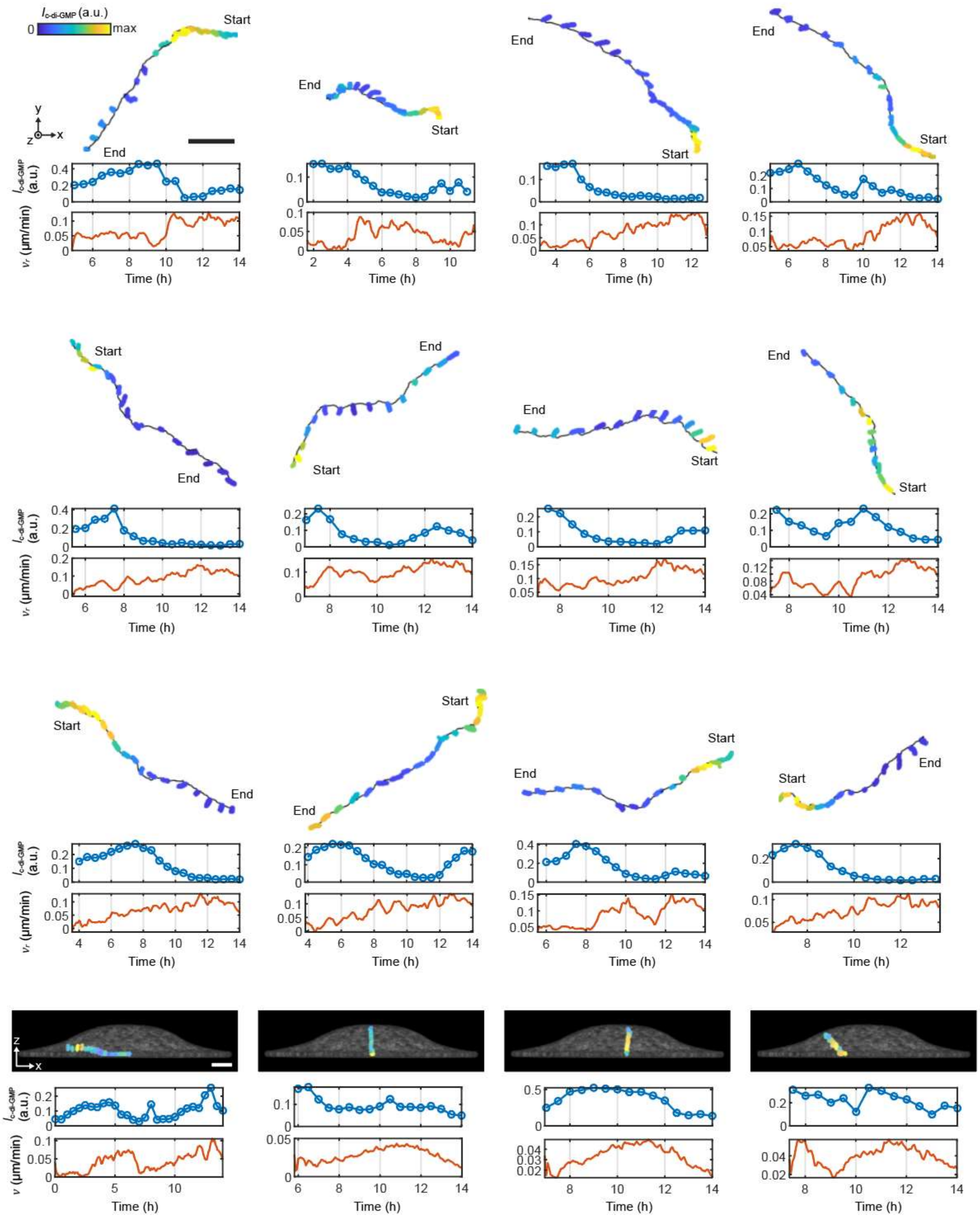

**Fig. S4. Example single-lineage trajectories in growing *V. cholerae* biofilms, related to Figure 4.** Single-lineage trajectories in growing WT *V. cholerae* biofilms, showing the segmented cells (*top*), $I_{\text{c-di-GMP}}$ vs. time (*middle*), and velocity vs. time (*bottom*) along each trajectory. Segmented cells are colored by $I_{\text{c-di-GMP}}$ based on the maximum $I_{\text{c-di-GMP}}$ value along each trajectory. The top three rows show trajectories within the bottom layer of the biofilms, and the bottom row shows trajectories that start from but subsequently move away from the glass substratum. The magnitude of the radial in-plane velocity $v_r$ is shown in the top three rows, while the magnitude of the 3D velocity is shown in the bottom row. In the bottom row, the translucent pattern shows the biofilm at the final timepoint of the plotted trajectory. Scale bars: 10 μm.

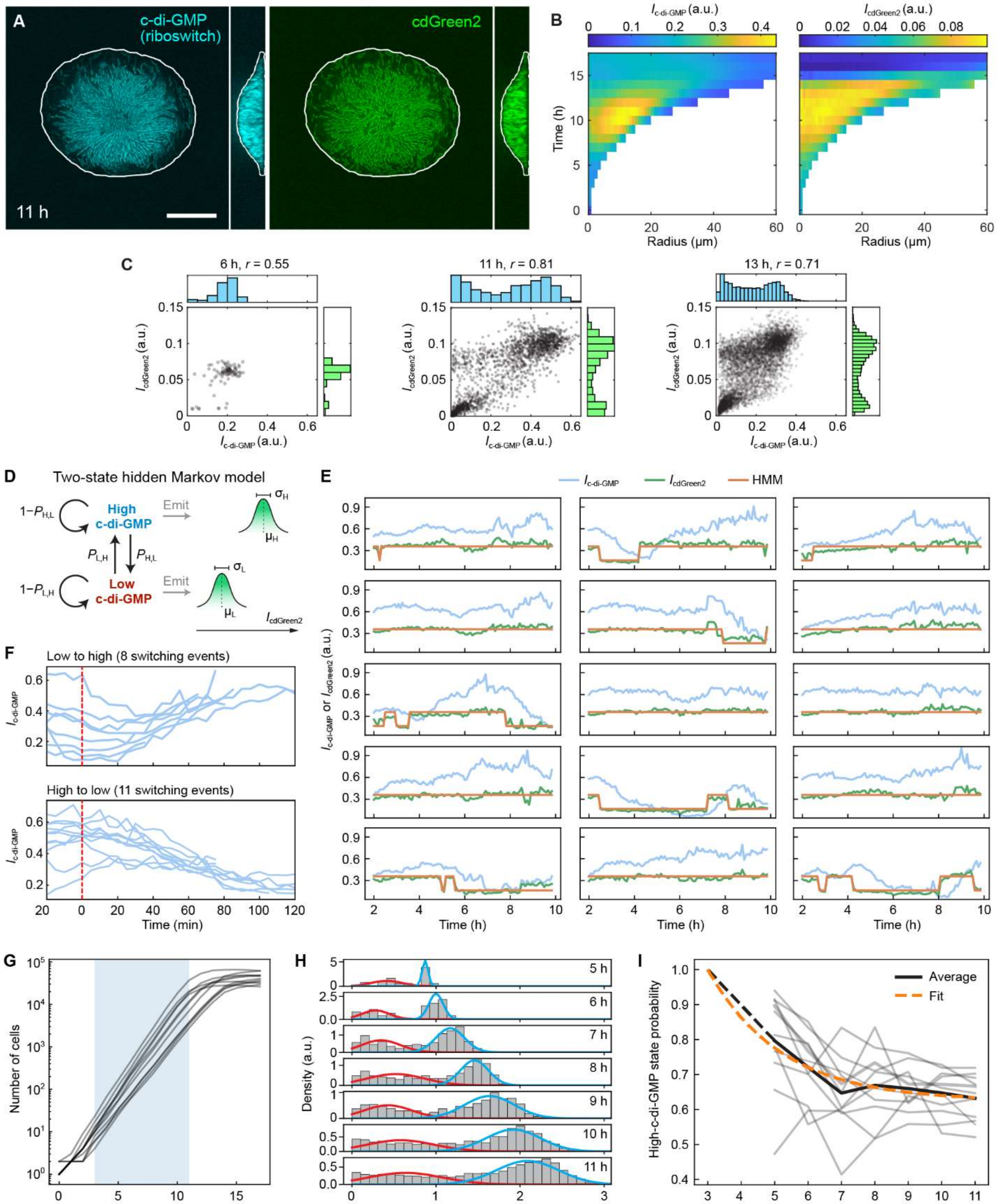

**Fig. S5. Comparison between the riboswitch-based c-di-GMP biosensor and cdGreen2, and determining switching rates of c-di-GMP phenotypes, related to Figure 5.**

(**A**) Confocal images of the bottom layer and vertical cross-section of a growing WT biofilm expressing both the riboswitch-based c-di-GMP biosensor (*left*) and cdGreen2 (*right*) at 11 h. Scale bar: 20 μm.

(**B**) Time vs. radius heatmaps of locally averaged $I_{\text{c-di-GMP}}$ (*left*) and $I_{\text{cdGreen2}}$ (*right*) of cells in the bottom layer of 10 WT biofilms.
(**C**) Correlation between $I_{\text{c-di-GMP}}$ and $I_{\text{cdGreen2}}$ of the biofilm shown in (A) throughout biofilm growth, as quantified by Pearson's correlation coefficient ($r$).
(**D**) Schematic of switching between low- and high-c-di-GMP phenotypes according to the two-state hidden Markov model (HMM; *Methods*). $P_{\text{L,H}}$ and $P_{\text{H,L}}$ are the transition probabilities at each discrete time step (5 min), and $\mu_{\text{L}}$ ($\mu_{\text{H}}$) and $\sigma_{\text{L}}$ ($\sigma_{\text{H}}$) are the mean and standard deviation of the Gaussian emission distribution corresponding to $I_{\text{cdGreen2}}$ for the low- (high-) c-di-GMP state.
(**E**) 15 additional example single-lineage traces of $I_{\text{c-di-GMP}}$ (cyan) and $I_{\text{cdGreen2}}$ (green) from 2 to 10 h, obtained from Δ*vpsL* colonies. Shown in orange are corresponding trajectories of predicted c-di-GMP phenotypes (low or high) according to the HMM, which was trained on a total of 51 single-lineage $I_{\text{cdGreen2}}$ traces.
(**F**) Collections of $I_{\text{c-di-GMP}}$ traces in lineages predicted to have undergone a low-to-high-c-di-GMP switch (*top*) or a high-to-low-c-di-GMP switch (*bottom*) according to the HMM based on $I_{\text{cdGreen2}}$, with no other switching events having occurred in the preceding 20 minutes or subsequent 40 minutes. Trajectories have been aligned so that these switching events occur at time zero (red). The realigned trajectories show that there is good agreement between transitions predicted by the HMM based on $I_{\text{cdGreen2}}$ and the change in $I_{\text{c-di-GMP}}$; they also show that there is a general delay in the response of $I_{\text{c-di-GMP}}$ to switching in $I_{\text{cdGreen2}}$, due to the requirement for transcription, translation, FP maturation, and/or dilution of the riboswitch-based c-di-GMP biosensor (but not cdGreen2) in response to changes in c-di-GMP level.
(**G**) Numbers of cells over time in 11 WT biofilms across three biological replicates. Fitting these curves between 3 and 11 h (blue) to an exponential growth function yielded a doubling time estimate of 48.6 ± 3.23 minutes (mean ± s.d.).
(**H** and **I**) Estimation of c-di-GMP phenotype lifetimes from population dynamics using Gaussian mixture modeling (*Methods*).
(**H**) Histograms of $I_{\text{c-di-GMP}}$ across a single WT biofilm from 5 to 11 h. A two-component Gaussian mixture model was fit to each $I_{\text{c-di-GMP}}$ distribution to estimate the proportions of low- and high-c-di-GMP cells (red and cyan, respectively).
(**I**) Proportions of high-c-di-GMP cells, estimated via Gaussian mixture modeling of $I_{\text{c-di-GMP}}$ distributions, in 11 WT biofilms from 5 to 11 h (grey). The average high-c-di-GMP proportion at each timepoint is shown in black; fitting this curve, with the additional assumption that biofilms are uniformly high-c-di-GMP at 3 h (dashed black line; *Methods*), to an exponential decay function representing the probability of the high-c-di-GMP state in the two-state model (orange; Figure 5A), yields estimates of $\tau_{\text{L}} \approx 3.55$ h and $\tau_{\text{H}} \approx 5.88$ h, which are somewhat longer than the trajectory-based estimates ($\tau_{\text{L}} \approx 2.14$ h and $\tau_{\text{H}} \approx 3.91$ h), but roughly agree in the ratio (Figure 5D).

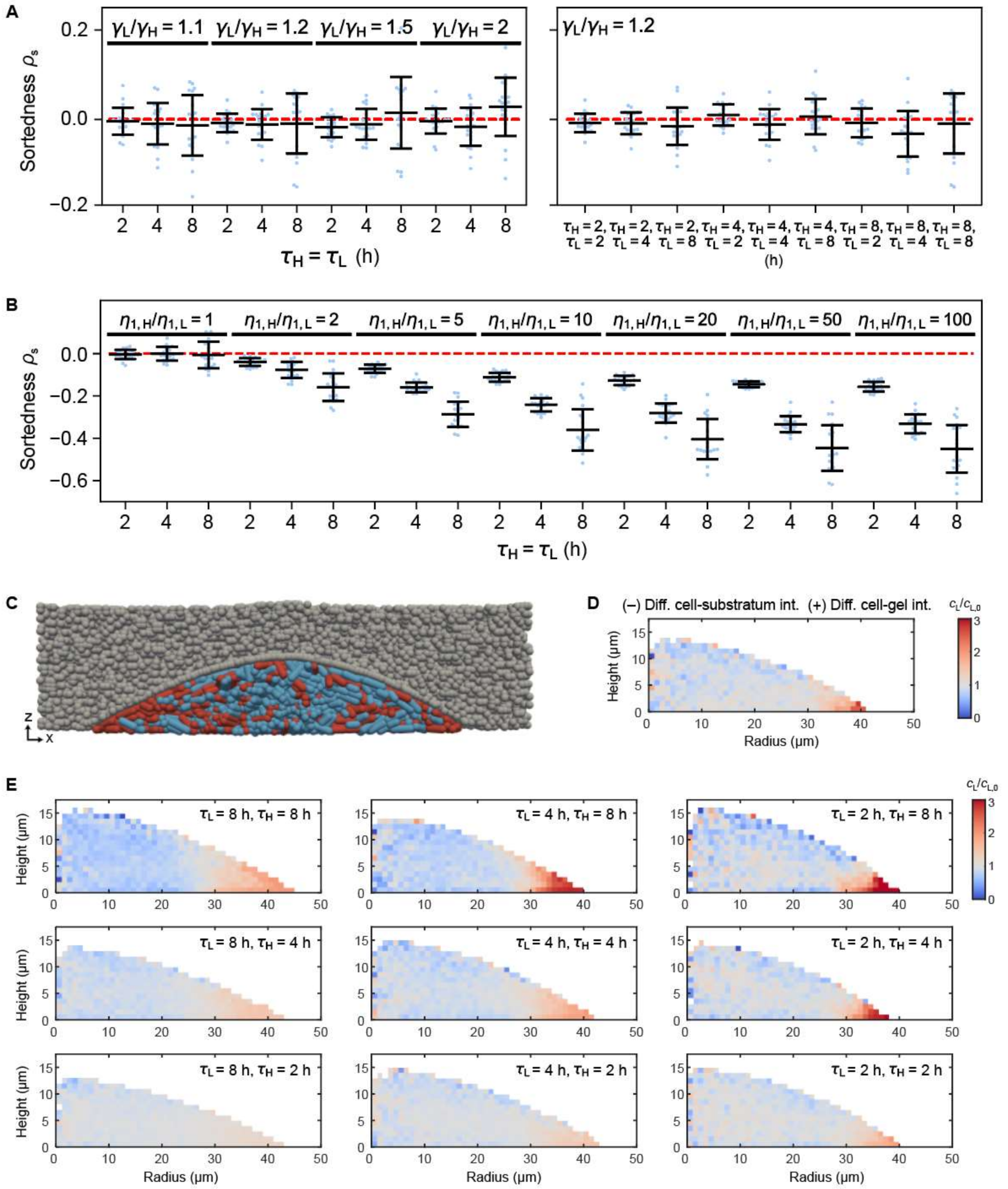

**Fig. S6. c-di-GMP-based sorting is reproduced by differential drag in 2D and 3D ABMs, related to Figure 5.**

(**A**) Sortedness quantification of simulation outcomes from 2D ABMs with different growth-rate ratios ($\gamma_L/\gamma_H$; Document S1) and mean state lifetimes ($\tau_L$ and $\tau_H$). Sortedness was quantified at 5000 cells in each simulation.

(**B**) Sortedness quantification of simulation outcomes from 2D ABMs with different generalized drag coefficient ratios (approximated as $\eta_{1,H}/\eta_{1,L}$; Document S1) and mean state lifetimes ($\tau_L$ and $\tau_H$). Sortedness was quantified at 5000 cells in each simulation.

(**C**) In 3D ABMs, cells with low- (red) and high-c-di-GMP (cyan) phenotypes were modeled to grow within the interstitial space between a rigid substratum and a hydrogel, where the latter was modeled as a connected network of spherical particles (grey).

(**D**) Distribution of c-di-GMP phenotypes, shown as the local relative fraction of low-c-di-GMP cells ($c_\mathrm{L}/c_{\mathrm{L},0}$), in the presence of differential interactions at the biofilm-gel interface but not along the substratum in 3D ABMs. $c_{\mathrm{L},0} \equiv \tau_\mathrm{H}^{-1}/(\tau_\mathrm{L}^{-1} + \tau_\mathrm{H}^{-1})$ is the theoretical steady-state low-c-di-GMP cell fraction across the population. Data are shown as $(r, z)$ projections averaged over 10 simulated biofilms ($\tau_\mathrm{L} = 2$ h and $\tau_\mathrm{H} = 4$ h).
(**E**) Distribution of c-di-GMP phenotypes, shown as the local relative fraction of low-c-di-GMP cells ($c_\mathrm{L}/c_{\mathrm{L},0}$), for various combinations of mean state lifetimes, $\tau_\mathrm{L}$ and $\tau_\mathrm{H}$, in 3D ABMs. Data are shown as $(r, z)$ projections, each averaged over 10 simulated biofilms. Here, both differential cell-substratum and cell-gel interactions are present.

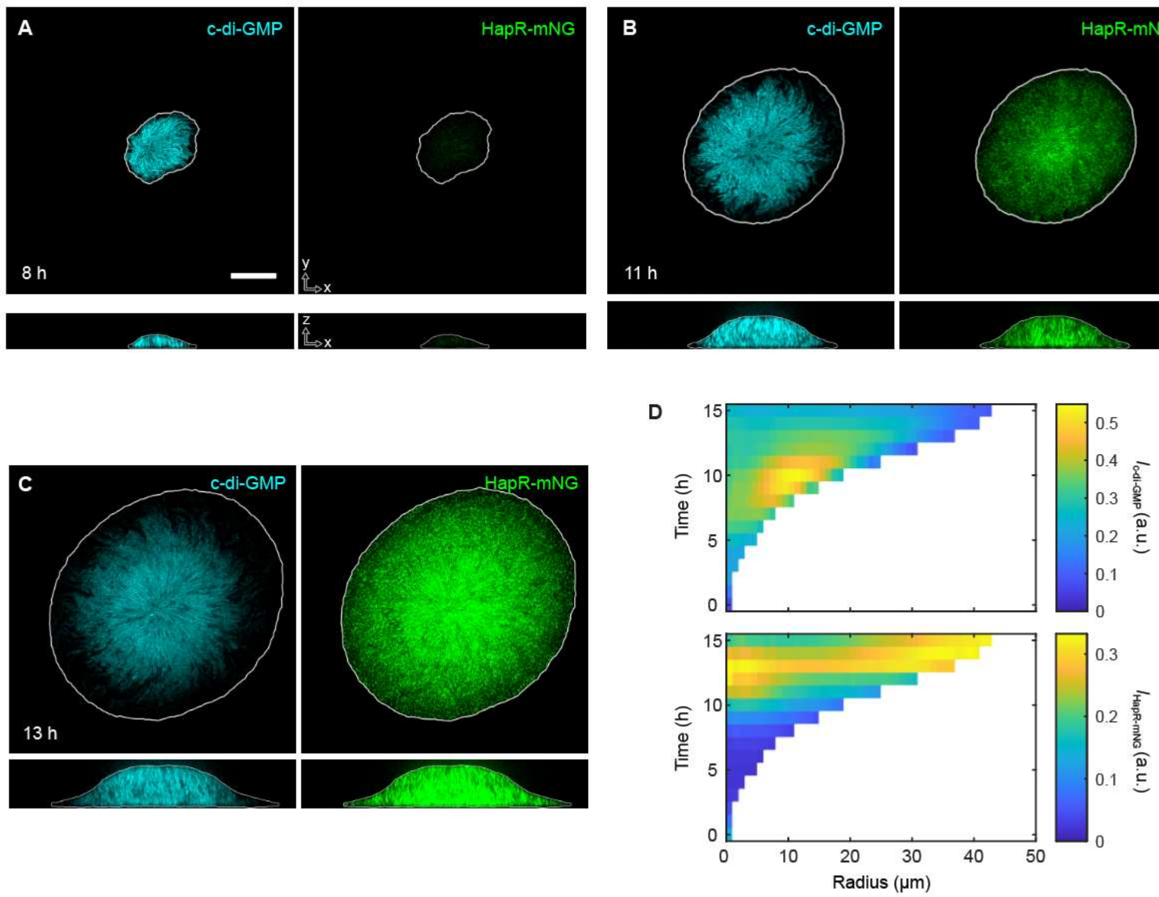

**Fig. S7. Spatiotemporal correlation between HapR and [c-di-GMP] in *V. cholerae* biofilms, related to Figure 6.** (**A–C**) Confocal images of the bottom layer and vertical cross-section of a WT biofilm expressing both the riboswitch-based c-di-GMP biosensor (*left*) and HapR-mNG (*right*) at 8 h (A), 11 h (B), and 13 h (C) during biofilm growth. Scale bar: 20 μm.

(**D**) Time vs. radius heatmaps of locally averaged $I_{\text{c-di-GMP}}$ (*top*) and $I_{\text{HapR-mNG}}$ (*bottom*) of cells in the bottom layer of WT biofilms. Data were obtained from 20 biofilms. $I_{\text{HapR-mNG}}$ increases notably around 8 h and develops higher intensity at the center, until becoming uniformly bright in the bottom layer at around 13 h; at even later times, $I_{\text{HapR-mNG}}$ decreases, especially at the center, for reasons yet unclear.

#### Table S1. *V. cholerae* strains used in this study

| Strain | Genotype | Source |
|---|---|---|
| JY016 | *Vibrio cholerae* O1 biovar El Tor strain C6706 str2 | |
| JST162 | ΔVC1807::P$_{tac}$-mScarlet-I-L3S2P21-P$_{tac}$-*Bc3-Bc4*-RBS-mNeonGreen(ASV), Spec$^{R}$ | This study |
| JST215 | Δ*rbmA* ΔVC1807::P$_{tac}$-mScarlet-I-L3S2P21-P$_{tac}$-*Bc3-Bc4*-RBS-mNeonGreen(ASV), Spec$^{R}$ | This study |
| JST208 | Δ*bap1* Δ*rbmC* ΔVC1807::P$_{tac}$-mScarlet-I-L3S2P21-P$_{tac}$-*Bc3-Bc4*-RBS-mNeonGreen(ASV), Spec$^{R}$ | This study |
| JST209 | Δ*rbmA* Δ*bap1* Δ*rbmC* ΔVC1807::P$_{tac}$-mScarlet-I-L3S2P21-P$_{tac}$-*Bc3-Bc4*-RBS-mNeonGreen(ASV), Spec$^{R}$ | This study |
| JST203 | Δ*vpsL* ΔVC1807::P$_{tac}$-mScarlet-I-L3S2P21-P$_{tac}$-*Bc3-Bc4*-RBS-mNeonGreen(ASV), Spec$^{R}$ | This study |
| JST257 | ΔVC1807::P$_{tac}$-mScarlet-I-L3S2P21-P$_{tac}$-*Bc3-Bc4*-RBS-SCFP3A(ASV), Spec$^{R}$ ΔVC0501::L3S2P21-P$_{vps\text{-}II}$-mNeonGreen(ASV)×2, Kan$^{R}$ | This study |
| JST269 | ΔVC1807::P$_{tac}$-mScarlet-I-L3S2P21-P$_{tac}$-*Bc3-Bc4*-RBS-SCFP3A(ASV), Spec$^{R}$ ΔVC0501::L3S2P21-P$_{rbmA}$-mNeonGreen(ASV)×2, Kan$^{R}$ | This study |
| JST309 | *rbmA*-3×FLAG ΔVC1807::P$_{tac}$-SCFP3A, Spec$^{R}$ ΔVC0501::L3S2P21-P$_{rbmA}$-2×mNeonGreen(ASV), Kan$^{R}$ | This study |
| JST315 | *rbmA*-TC ΔVC1807::P$_{tac}$-SCFP3A, Spec$^{R}$ ΔVC0501::L3S2P21-P$_{rbmA}$-2×mNeonGreen(ASV), Kan$^{R}$ | This study |
| JST268 | ΔVC1807::P$_{tac}$-mScarlet-I-L3S2P21-P$_{tac}$-*Bc3-Bc4*-RBS-SCFP3A(ASV), Spec$^{R}$ Δ*lacIZ*:: P$_{tac}$-mNeonGreen-μNS | This study |
| JST285 | Δ*vpsL* ΔVC1807::P$_{tac}$-mScarlet-I-L3S2P21-P$_{tac}$-*Bc3-Bc4*-RBS-SCFP3A(ASV), Spec$^{R}$ Δ*lacIZ*:: P$_{tac}$-mNeonGreen-μNS | This study |
| JST276 | ΔVC1807::P$_{tac}$-mScarlet-I-L3S2P21-P$_{tac}$-*Bc3-Bc4*-RBS-mNeonGreen(ASV), Spec$^{R}$ pJST35 | This study |
| JST278 | *vpvC*$^{W240R}$ ΔVC1807::P$_{tac}$-mScarlet-I-L3S2P21-P$_{tac}$-*Bc3-Bc4*-RBS-mNeonGreen(ASV), Spec$^{R}$ pJST37 | This study |
| JST338 | ΔVC1807::P$_{tac}$-mScarlet-I-L3S2P21-P$_{tac}$-*Bc3-Bc4*-RBS-SCFP3A(ASV), Spec$^{R}$ ΔVC0501::P$_{tac}$-cdGreen2, Kan$^{R}$ | This study |
| JST339 | Δ*vpsL* ΔVC1807::P$_{tac}$-mScarlet-I-L3S2P21-P$_{tac}$-*Bc3-Bc4*-RBS-SCFP3A(ASV), Spec$^{R}$ ΔVC0501::P$_{tac}$-cdGreen2, Kan$^{R}$ | This study |
| JST363 | Δ*pomA* Δ*bap1* Δ*rbmC* ΔVC1807::P$_{tac}$-mScarlet-I-L3S2P21-P$_{tac}$-*Bc3-Bc4*-RBS-mNeonGreen(ASV), Spec$^{R}$ | This study |
| JST271 | Δ*hapR*::*hapR*-mNeonGreen ΔVC1807::P$_{tac}$-mScarlet-I-L3S2P21-P$_{tac}$-*Bc3-Bc4*-RBS-SCFP3A(ASV), Spec$^{R}$ | This study |
| JST272 | Δ*hapR* ΔVC1807::P$_{tac}$-mScarlet-I-L3S2P21-P$_{tac}$-*Bc3-Bc4*-RBS-mNeonGreen(ASV), Spec$^{R}$ | This study |
| JST274 | ΔVC1807::P$_{tac}$-mScarlet-I, Spec$^{R}$ ΔVC0501::P$_{tac}$-HYlight, Kan$^{R}$ | |
| JST204 | Δ*pomA* ΔVC1807::P$_{tac}$-mScarlet-I-L3S2P21-P$_{tac}$-*Bc3-Bc4*-RBS-mNeonGreen(ASV), Spec$^{R}$ | This study |
| JST348 | ΔVC1807::P$_{tac}$-mScarlet-I, Spec$^{R}$ ΔVC0501::P$_{BAD}$-VC1086, Kan$^{R}$ | This study |
| JST349 | ΔVC1807::P$_{tac}$-SCFP3A, Spec$^{R}$ ΔVC0501::P$_{BAD}$-*qrgB*, Kan$^{R}$ | This study |
| JST352 | ΔVC1807::P$_{tac}$-mNeonGreen, Spec$^{R}$ ΔVC0501::P$_{BAD}$-VC1086$^{*}$ (inactive), Kan$^{R}$ | This study |
| JST264 | ΔVC1807::P$_{tac}$-mScarlet-I-L3S2P21-P$_{tac}$-mNeonGreen(ASV), Spec$^{R}$ | This study |
| JST290 | ΔVC1807::P$_{tac}$-mScarlet-I-L3S2P21-P$_{tac}$-SCFP3A(ASV), Spec$^{R}$ ΔVC0501::P$_{tac}$-mNeonGreen(ASV), Kan$^{R}$ | This study |
| JST326 | Δ*vpsL* ΔVC1807::P$_{tac}$-mScarlet-I-L3S2P21-P$_{tac}$-*Bc3-Bc4*-RBS-mNeonGreen(ASV), Spec$^{R}$ ΔVC0501::P$_{BAD}$-*vpvC*$^{W240R}$, Kan$^{R}$ | This study |
| JST301 | Δ*vpsL* ΔVC1807::P$_{tac}$-mScarlet-I-L3S2P21-P$_{tac}$-*Bc3-Bc4*-RBS-mNeonGreen(ASV), Spec$^{R}$ ΔVC0501::P$_{BAD}$-VC1086, Kan$^{R}$ | This study |
| **Plasmid** | | |
| pJST35 | Kan$^{R}$, P$_{tac}$-*qrgB*, P$_{lacI}$-SCFP3A (pEVS141 backbone) | This study |
| pJST37 | Kan$^{R}$, P$_{tac}$-VC1086, P$_{lacI}$-SCFP3A (pEVS141 backbone) | This study |

#### Table S2. Oligonucleotides used in this study

| Number | Oligonucleotide sequence (5' to 3') | Description |
|---|---|---|
| pJY121 | tttaaaggggatcagtgaccg | Forward primer annealing to 3 kb upstream of VC1807 |
| pJY122 | caattttgcttttggaccatccc | Reverse primer annealing to 3 kb downstream of VC1807 |
| pJY135 | ggccggcactttgattacaatc | Forward primer annealing to 2.7 kb upstream of VC1807 |
| pJY136 | gtctatatcagagcgcttaaagagcg | Reverse primer annealing to 2.7 kb downstream of VC1807 |
| oJST107 | cgttaccggatgtttacgcttagg | Forward primer annealing to 2.3 kb upstream of VC0501 |
| oJST108 | atcgcgtcttagtatgcgagac | Reverse primer annealing to 2.3 kb downstream of VC0501 |
| oJST114 | gcgacttggcttgatgtcaatc | Forward primer annealing to 2 kb upstream of VC0501 |
| oJST115 | atgttatctattcggacagcgattactgtg | Reverse primer annealing to 2 kb downstream of VC0501 |
| oJST075 | taagaattcctcggtaccaaattccagaaaagaggcctcccgaaaggggggccttttt cgttttggtccactagtagcggccgctgcag | L3S2P21 transcriptional terminator (double-stranded ultramer) |
| oJST078 | cttttctggaatttggtaccgaggaattcttattatttgtacaactcgtccatcccc | Reverse primer annealing to the 3' end of the mScarlet-I sequence with overlap to the downstream L3S2P21 region |
| oJST084 | ggtccactagtagcggccgctgcagtgcaccaatgcttctggcgtc | Forward primer annealing to the 5' end of $P_{tac}$ with overlap to the upstream L3S2P21 sequence |
| oJST040 | aattgaattcctaggcctgtcgagg | Reverse primer annealing to the 3' end of $P_{tac}$ |
| oJST041 | cctcgacaggcctaggaattcaatttaaggatccacgataaataaatacctatttttggc | Forward primer annealing to the 5' end of *Bc3-Bc4* riboswitch sequence with overlap to the upstream $P_{tac}$ sequence |
| oJST031 | atctgttttgtcgacaaaataaaaagcctg | Reverse primer annealing to the 3' end of *Bc3-Bc4* riboswitch sequence |
| oJST032 | caggctttttattttgtcgacaaaacagataggaggtaattaagcatggtatcgaagg | Forward primer annealing to the ribosome binding site and the 5' end of the mNeonGreen sequence with overlap to the upstream *Bc3-Bc4* sequence |
| oJST144 | ctggaatttggtaccgaggaattcttattgaagtctcgagatgcgatatctattgatg | Reverse primer annealing to the 3' end of VC0501 upstream homologous region with overlap to the downstream L3S2P21 sequence |
| oJST076 | taagaattcctcggtaccaaattccagaaaag | Former primer annealing to the 5' end of L3S2P21 |
| oJST077 | ctgcagcggccgctactag | Reverse primer annealing to the 3' end of L3S2P21 |
| oJST079 | cactagtagcggccgctgcagtttgattaacctattaaccatcataaaagtaaactaaag | Forward primer annealing to the 5' end of $P_{vps\text{-II}}$ with overlap to the upstream L3S2P21 sequence |
| oJST224 | ccttcgataccatgcttaattacctcctaaccgatgtaagatttcccgaagaataattg | Reverse primer annealing to the 3' end of $P_{vps\text{-II}}$ with overlap to the downstream ribosome binding site and mNeonGreen sequence |
| oJST039 | aggaggtaattaagcatggtatcgaagg | Forward primer annealing to the ribosome binding site and the 5' end of the mNeonGreen sequence |
| oJST092 | cactagtagcggccgctgcaggttacaagaacccggaagaatgtgg | Forward primer annealing to the 5' end of PrbmA with overlap to the upstream L3S2P21 sequence |
| oJST093 | ccttcgataccatgcttaattacctcctccatttgtttttacaactggcgctaag | Reverse primer annealing to the 3' end of $P_{rbmA}$ with overlap to the downstream ribosome binding site and mNeonGreen sequence |
| pJY188 | tggtgttgtccgggctgctgtaagactttttaaatttacctagtcacttagtcgtatgta | Former primer introducing the tetracysteine motif (HRWCCPGCCKTF) into RbmA |

| | | |
|---|---|---|
| pJY189 | cttacagcagcccggacaacaccaacgatgtttttttaccactgtcattgactgttccac | Reverse primer introducing the tetracysteine motif (HRWCCPGCCKTF) into RbmA |
| oJST232 | gcctaggaattcaattaggaggtaattaagcatgaattcagagccgccgc | Former primer annealing to the 5' end of cdGreen2 sequence with overlap to the upstream $P_{tac}$ and the ribosome binding site sequences |
| oJST310 | gtcgacggatccccggaattcagttctcctcgtgggcg | Reverse primer annealing to the 3' end of cdGreen2 sequence with overlap to the downstream antibiotic-resistance sequence |
| oJST230 | gttgtcataattggtaacgaatcagacaattttgaagtctcgagatgcgatatctattg | Reverse primer annealing to the 3' end of VC0501 upstream homologous region with overlap to the downstream $P_{BAD}$ sequence |
| oJST229 | aattgtctgattcgttaccaattatgacaac | Forward primer annealing to the 5' end of $P_{BAD}$ sequence |
| oJST206 | tttagacctccttactggtactgcggccgcggagaaacag | Reverse primer annealing to the 3' end of $P_{BAD}$ sequence |
| oJST205 | cgcagtaccagtaaggaggtctaaaatgcaaagcaaccgtg | Former primer annealing to the 5' end of VC1086 sequence with overlap to the upstream $P_{BAD}$ sequence |
| oJST236 | cgcagtaccagtaaggaggtctaaagtgatcggtattgattgtgaactcaatataaaca g | Former primer annealing to the 5' end of *qrgB* sequence with overlap to the upstream $P_{BAD}$ sequence |
| oJST231 | gtcgacggatccccggaatgcttgctcaatcaatcaccggatc | Reverse primer annealing to the sequence after the 3' end of VC1086 or *qrgB* on pYS249 plasmid backbone with overlap to the downstream antibiotic-resistance sequence |
| oJST307 | cgcagtaccagtaaggaggtctaaaatgactgatcaaacgcgaacttcg | Former primer annealing to the 5' end of *vpvC* sequence with overlap to the upstream $P_{BAD}$ sequence |
| oJST308 | gtcgacggatccccggaatctatctgaactgatcctgcttgagttc | Reverse primer annealing to the 3' end of *vpvC* sequence with overlap to the downstream antibiotic-resistance sequence |
| oJST175 | gcgcaacgcaattaatgtaagttagc | Forward primer annealing to pEVS141 plasmid backbone |
| oJST176 | tactggtacttcactgcccgctttccag | Reverse primer annealing to the 3' end of *lacI* sequence |
| oJST258 | ctggaaagcgggcagtgaagtaccagtaaggaggtaattaagcatgtctaagggtg | Forward primer annealing to the ribosome binding site and the 5' end of SCFP3A sequence with overlap to the upstream *lacI* sequence |
| oJST259 | cgcgctaacttacattaattgcgttgcgcttacttgtacaattcgtccatacccaaagtg | Reverse primer annealing to the 3' end of SCFP3A sequence with overlap to the downstream sequence of pEVS141 plasmid backbone |
| oJST094 | cgtaattctcgtcattagcagcgggacgcttatataattcatccatccccataacgtctg | Reverse primer introducing the ASV tag (AANDENYAASV) at the C-terminus of mNeonGreen |
| oJST142 | cgtaattctcgtcattagcagcgggacgcttgtacaattcgtccatacccaaagtgatac | Reverse primer introducing the ASV tag (AANDENYAASV) at the C-terminus of SCFP3A |
| oJST106 | gtcccgctgctaatgacgagaattacgcggcatcggtttaaattccggggatccgtcg ac | Former primer introducing the ASV tag (AANDENYAASV) at the C-terminus of mNeonGreen and SCFP3A |
