## Supplementary material for "Single-cell imaging reveals spontaneous phenotypic sorting and bet-hedging in developing biofilms": Document S1

**Document S1: Description of agent-based models.**

Here, we describe the agent-based models (ABMs) that we implemented in this study. The 3D ABM implemented in this paper is an extension of a previously developed ABM, whose details can be found in the corresponding papers[1–4]; the 2D ABM was developed by appropriately simplifying the 3D ABM. As such, we begin by describing the ways in which we extended the 3D ABM from prior work, focusing on the two-state model of phenotypic switching (**Section 1**); and how the 2D ABM arises as a simplified version of the 3D ABM (**Section 2**). We then discuss our method for imposing differential phenotypes on the low- and high-c-di-GMP cells (**Section 3**) and our implementation of the growth void as a proxy for verticalization in the 2D ABM (**Section 4**), and finally enumerate the parameter values used in our simulations (**Tables 1–3**).

### 1. 3D ABM.

The 3D ABM implemented in this study builds upon a previously developed 3D ABM for a *Vibrio cholerae* biofilm growing within the interstitial space between a glass substratum and an overlaid agarose gel[1–4]. Briefly, this ABM describes each cell as an elongating, matrix-encased spherocylinder that divides upon reaching a critical length; describes the agarose gel as a coarse-grained, mechanical network of spherical particles; and includes mechanical forces arising from cell-cell repulsion, cell-substratum repulsion, cell-substratum adhesion, cell-gel repulsion, and cell-gel adhesion, as well as viscous forces on each cell owing to ambient viscosity within the bulk of the biofilm and cell-substratum friction. The three updates we have introduced in this work are: (1) a modified model of cell-cell repulsion, (2) a model of cell-gel friction, and (3) phenotypic switching according to the two-state model (Figure 5A), which we describe below.

***Preliminaries.*** In what follows, we let $\mathbf{r}_i$ be the center of cell $i$; let $\hat{\mathbf{n}}_i$ be the unit vector defining the orientation of cell $i$; let $l_i$ be the length of cell $i$; and let $R$ and $R_{\text{cell}}$ be the cell radius with and without the matrix, respectively. Similarly, if $j$ is the index for a gel particle, we denote by $\mathbf{r}_j$ the center of gel particle $j$, and let $R_{\text{gel}}$ be the gel particle radius, which is the same for all gel particles.

For any two bodies $i$ and $j$, we denote by $\mathbf{d}_{i,j}$ the shortest vector that runs from the center or centerline of body $i$ to that of body $j$; we denote by $\hat{\mathbf{d}}_{i,j}$ the corresponding unit vector; and we denote by $\delta_{i,j}$ the theoretical overlap, which we define as

$$\delta_{i,j} = \left(R_i + R_j - \left\|\mathbf{d}_{i,j}\right\|\right) \Theta\left(R_i + R_j - \left\|\mathbf{d}_{i,j}\right\|\right),$$

where $R_k$ is the radius of body $k$ ($R_k = R$ if body $k$ is a cell, and $R_k = R_{\text{gel}}$ if body $k$ is a gel particle), and $\Theta(\cdot)$ is the Heaviside step function.

Finally, we write $\mathbf{v}_i = d\mathbf{r}_i/dt$ for the velocity of body $i$ at its center. If body $i$ is a cell, then we write $\mathbf{v}_i(s) = d\mathbf{r}_i/dt + s \cdot d\hat{\mathbf{n}}_i/dt$ for the cell's velocity at the point $\mathbf{r}_i + s\hat{\mathbf{n}}_i$, for $s \in [-l_i/2, l_i/2]$, along the cell's centerline, and we write $\boldsymbol{\omega}_i$ for the angular velocity of the cell.

***Cell-cell repulsion.*** Following the previous iteration of this ABM[3], we model each cell as a stiff spherocylinder with elastic modulus $E_{\text{cell}}$ that is encased in a uniform shell of extracellular matrix, which has a lower elastic modulus $E_0$. As such, we modeled the cell-cell repulsion force on cell $i$ due to cell $j$ as a Hertzian force[5] that accounts for the two elastic moduli, as

$$\mathbf{F}_{\text{ccr},i,j} = \begin{cases} \mathbf{0} & \text{if } d > 2R \\ -\frac{5}{2} E_0 R^{1/2} (2R - d)^{3/2} \hat{\mathbf{d}}_{i,j} & \text{if } 2R_{\text{cell}} < d \leq 2R \\ -\frac{5}{2} \left(E_0 R^{1/2} (2R - 2R_{\text{cell}})^{3/2} + E_{\text{cell}} (R_{\text{cell}})^{1/2} (2R_{\text{cell}} - d)^{3/2}\right) \hat{\mathbf{d}}_{i,j} & \text{if } d \leq 2R_{\text{cell}}, \end{cases} \qquad 1$$

where $d = \left\|\mathbf{d}_{i,j}\right\|$. This force differs slightly from that in ref. [3] in how the soft-to-hard-contact transition is implemented; we believe that this definition handles this transition more appropriately. The corresponding moment on cell $i$ due to cell $j$ is given by

$$\mathbf{M}_{\mathrm{ccr},i,j} = s^*\hat{\mathbf{n}}_i \times \mathbf{F}_{\mathrm{ccr},i,j},$$

where $s^* \in [-l_i/2, l_i/2]$ is the centerline coordinate along cell $i$ at which the contact occurs (i.e., the tail of the distance vector, $\mathbf{d}_{i,j}$, is positioned at the point $\mathbf{r}_i + s^*\hat{\mathbf{n}}_i$).

***Cell-gel friction.*** To model the tangential friction forces that arise from adhesive contacts between each cell and the agarose gel, we used a simple viscous force that depends linearly on the relative tangential velocity between the contacting cell and gel particle. Specifically, the friction force on cell $i$ due to gel particle $j$ is given by:

$$\mathbf{F}_{\mathrm{cgf},i,j} = -\eta_{\mathrm{cg}}\sqrt{\delta_{i,j}R_{\mathrm{eq}}}\mathbf{v}_{i,j}^{\mathrm{t}}, \qquad 2$$

where $\eta_{\mathrm{cg}}$ is a friction coefficient, $R_{\mathrm{eq}} = 2R_{\mathrm{gel}}R/(R + R_{\mathrm{gel}})$ is the equivalent radius of contact, and $\mathbf{v}_{i,j}^{\mathrm{t}}$ is the tangential component of the relative velocity, $\mathbf{v}_{i,j}^*$, at the contact point:

$$\mathbf{v}_{i,j}^{\mathrm{t}} = \mathbf{v}_{i,j}^* - (\mathbf{v}_{i,j}^* \cdot \hat{\mathbf{d}}_{i,j})\hat{\mathbf{d}}_{i,j},$$

where the relative velocity is given by

$$\mathbf{v}_{i,j}^* = \mathbf{v}_i - \left(R - \frac{\delta_{i,j}}{2}\right)\boldsymbol{\omega}_i \times (\mathbf{r}_i - \mathbf{r}_j) - \mathbf{v}_j.$$

This definition follows that of similar forces that are often included in discrete element method simulations of granular materials[6–12].

***Two-state model.*** We used the two-state model (Figure 5A) to model heterogeneity in c-di-GMP signaling among cells within a biofilm. Specifically, we assumed that each cell switches between two states, $\mathrm{L}$ (for low-c-di-GMP) and $\mathrm{H}$ (for high-c-di-GMP), according to a continuous-time Markov chain with mean state lifetimes $\tau_{\mathrm{L}}$ and $\tau_{\mathrm{H}}$, respectively. In particular, this means that, if $\sigma_i(t) \in \{\mathrm{L}, \mathrm{H}\}$ is the state of cell $i$ at time $t$, then the infinitesimal transition rate from, e.g., $\mathrm{L}$ to $\mathrm{H}$ is given by[13]

$$\lim_{\Delta t \to 0} \frac{\Pr(\sigma_i(t + \Delta t) = \mathrm{H} \mid \sigma_i(t) = \mathrm{L})}{\Delta t} = \frac{1}{\tau_{\mathrm{L}}}.$$

As such, we applied the following algorithm at each timestep, $[t, t + \Delta t)$, to determine $\sigma_i(t + \Delta t)$ from $\sigma_i(t)$ for each cell:

1. Sample a random number, $r$, from the uniform distribution on $[0,1]$.
2. Set $\sigma_i(t + \Delta t)$ as follows:
    a. If $\sigma_i(t) = \mathrm{L}$ and $r < \Delta t/\tau_{\mathrm{L}}$, then set $\sigma_i(t + \Delta t) = \mathrm{H}$ and sample new value(s) for the attribute(s) of cell $i$ accordingly.
    b. If $\sigma_i(t) = \mathrm{H}$ and $r < \Delta t/\tau_{\mathrm{H}}$, then set $\sigma_i(t + \Delta t) = \mathrm{L}$ and sample new value(s) for the attribute(s) of cell $i$ accordingly.
    c. Otherwise, set $\sigma_i(t + \Delta t) = \sigma_i(t)$.

In each simulation, the low- and high-c-di-GMP states were associated with different values (or distributions of values) for one or more phenotypic attributes of each cell. We enumerate these simulation assumptions in **Section 3**.

### 2. 2D ABM.

The 2D ABM was formulated from the 3D ABM by assuming that each cell is perfectly parallel to the glass substratum. We assumed that the combined effects of gel confinement were to maintain this 2D geometry, and thus omitted the cell-gel repulsion, adhesion, and friction forces from the model. Furthermore, we set the *z*-position of each cell to a value that minimizes the energetic contributions of cell-substratum repulsion and adhesion. The cell-substratum repulsion energy is given, for cell $i$, by[1]

$$E_{i,\mathrm{csr}} = E_0 R^{1/2} \int_{-l_i/2}^{l_i/2} \left( R^{-1/2} \left(1 - \left|\hat{n}_{i,z}\right|^2\right) \delta_i^2(s) + \frac{4}{3}\left|\hat{n}_{i,z}\right|^2 \delta_i^{3/2}(s) \right) \mathrm{d}s \,,$$

where $E_0$ is the elastic modulus of the matrix, $l_i$ is the cell length, and $\delta_i(s)$ is the theoretical cell-substratum overlap at the point $\mathbf{r}_i + s\hat{\mathbf{n}}_i$, for $s \in [-l_i/2, l_i/2]$, along the cell's centerline, which is given by

$$\delta_i(s) = \left(R - \left(r_{i,z} + s\hat{n}_{i,z}\right)\right) \Theta \left(R - \left(r_{i,z} + s\hat{n}_{i,z}\right)\right).$$

On the other hand, the cell-substratum adhesion energy is given by[1]

$$E_{i,\mathrm{csa}} = -\Sigma_0 \int_{-l_i/2}^{l_i/2} \left( R^{1/2} \left(1 - \left|\hat{n}_{i,z}\right|^2\right) \delta_i^{1/2}(s) + \pi R \left|\hat{n}_{i,z}\right|^2 \Theta\left(\delta_i(s)\right) \right) \mathrm{d}s \,, \quad 3$$

where $\Sigma_0$ is a constant adhesion energy density. Assuming that each cell is horizontal ($\hat{n}_{i,z} = 0$), differentiating $E_{i,\mathrm{csr}} + E_{i,\mathrm{csa}}$ with respect to $r_{i,z}$ and setting the derivative equal to zero yields

$$r_{i,z} = R - \left(\frac{\Sigma_0 R^{1/2}}{4E_0}\right)^{2/3}.$$

Thus, setting the *z*-position of each cell to this value was sufficient to omit all cell-substratum adhesion and repulsion forces from the model as well. The resulting equations of motion, formulated in terms of Lagrangian mechanics[1], read as follows (see also, e.g., ref. [14]):

$$\left(\eta_{0,i} l_i + \frac{\eta_{1,i} a_i l_i}{R}\right) \frac{d\mathbf{r}_i}{dt} = -\frac{\partial E_{\mathrm{ccr},i}}{\partial \mathbf{r}_i} \quad 4$$

$$\left(\frac{\eta_{0,i} l_i^3}{12} + \frac{\eta_{1,i} a_i l_i^3}{12R}\right) \frac{d\hat{\mathbf{n}}_i}{dt} = -\frac{\partial E_{\mathrm{ccr},i}}{\partial \hat{\mathbf{n}}_i} + \lambda \frac{\partial}{\partial \hat{\mathbf{n}}_i} \left(\hat{\mathbf{n}}_i \cdot \hat{\mathbf{n}}_i - 1\right), \quad 5$$

where $\eta_{0,i}$ and $\eta_{1,i}$ are the dynamic viscosity and cell-substratum friction coefficient experienced by the cell, respectively; $E_{\mathrm{ccr},i}$ is the total cell-cell repulsion energy, $E_{\mathrm{ccr},i} = \sum_{j \neq i} E_{\mathrm{ccr},i,j}$, where (see also Eqn. 1)

$$E_{\mathrm{ccr},i,j} = \begin{cases} 0 & \text{if } d > 2R \\ E_0 R^{1/2} (2R - d)^{5/2} & \text{if } 2R_{\mathrm{cell}} < d \leq 2R \\ \frac{5}{2} E_0 R^{1/2} (2R - 2R_{\mathrm{cell}})^{3/2} (d - d_0) + E_{\mathrm{cell}} (R_{\mathrm{cell}})^{1/2} (2R_{\mathrm{cell}} - d)^{5/2} & \text{if } d \leq 2R_{\mathrm{cell}}, \end{cases}$$

with $d_0$ is a constant that ensures that the energy is continuous at $d = 2R_{\mathrm{cell}}$; $\lambda$ is a Lagrange multiplier; and $a_i$ is a cell-substratum contact area density, which is fixed as

$$a_i = R^{1/2} \delta_i^{1/2} = \left(\frac{\Sigma_0 R^2}{4E_0}\right)^{1/3}.$$

Note that the left-hand sides of Eqns. 4 and 5 can be written in terms of a generalized drag coefficient,

$$\eta_i = \eta_{0,i} + \frac{\eta_{1,i} a_i}{R}, \quad 6$$

which combines the effects of ambient viscosity ($\eta_{0,i}$) and cell-substratum friction ($\eta_{1,i}$), since every cell contacts the substratum to the same extent in 2D. This generalized drag coefficient is what we modulated to impose the differential drag mechanism, as described in **Section 3**.

**3. Imposing differential phenotypes between low- and high-c-di-GMP cells.**

For each ABM in this study, we assumed that the low- and high-c-di-GMP cells differ with respect to a phenotypic parameter, $\theta$, whose values we denote by $\theta_{\mathrm{L}}$ and $\theta_{\mathrm{H}}$, respectively. All other parameters were assumed to take the default values given in **Tables 1–3**.

***Differential growth rates (Figures 5E and S6A).*** In these simulations, the growth rate, $\gamma_i$, of each cell was assumed to switch between two normal distributions with means $\gamma_{\mathrm{H}}$ and $\gamma_{\mathrm{L}}$ and standard deviations $0.2\gamma_{\mathrm{H}}$ and $0.2\gamma_{\mathrm{L}}$, respectively. (The growth rate, $\gamma_i$, of cell $i$ determines the time-evolution of the cell's volume, $V_i$, as $dV_i/dt = \gamma_i V_i$; see ref. [1] for details.) We fixed $\gamma_H$ at the value previously estimated[1] for a constitutively biofilm-forming, or "rugose," *V. cholerae* strain[17] with an *rbmA* deletion, $\gamma_H = 3.12 \cdot 10^{-4}\ \mathrm{s}^{-1}$ (which is roughly $1.12\ \mathrm{h}^{-1}$), and set $\gamma_L = K\gamma_H$ for $K = 1.1, 1.2, 1.5, 2$.

***Differential drag in 2D (Figures 5F, 5G, and S6B).*** In the 2D ABMs, we imposed differential drag by modulating the generalized drag coefficient, $\eta_i$, in Eqn. 6. Now, since the default value of $\eta_{1,i}$ far exceeds that of $\eta_{0,i}$ ($\eta_{1,i} = 10^4\eta_{0,i}$, **Table 1**) and $R = 0.8\ \mu\mathrm{m}$ and $a_i \approx 0.16\ \mu\mathrm{m}$ are comparable in value (**Table 1**), we assumed that the effects of switching $\eta_i$ between two values could be captured by simply switching $\eta_{1,i}$ between two values, $\eta_{1,\mathrm{H}}$ and $\eta_{1,\mathrm{L}}$. Namely, we fixed $\eta_{1,\mathrm{H}}$ at the value previously estimated[1] for the Δ*rbmA* rugose *V. cholerae* strain, $\eta_{1,\mathrm{H}} = 2 \cdot 10^5\ \mathrm{Pa \cdot s}$ (which is roughly $720\ \mathrm{kg \cdot \mu m^{-1} \cdot hr^{-1}}$), and set $\eta_{1,\mathrm{L}} = \eta_{1,\mathrm{H}}/K$ for $K = 1, 2, 5, 10, 20, 50, 100$.

***Differential cell-substratum friction in 3D (Figures 5H and S6E).*** In the 3D ABMs, the cell-substratum friction force was assumed to follow the form[1–4],

$$\mathbf{F}_{i,\mathrm{csf}} = -\eta_{1,i} \int_{-l_i/2}^{l_i/2} \frac{a_i(s)}{R} \left(\mathbf{v}_i(s) - (\mathbf{v}_i(s) \cdot \hat{\mathbf{z}})\hat{\mathbf{z}}\right) \mathrm{d}s,$$

where $\eta_{1,i}$ is a cell-substratum friction coefficient; $\mathbf{v}_i(s) = d\mathbf{r}_i/dt + s \cdot d\hat{\mathbf{n}}_i/dt$ is the velocity of the cell at the point $\mathbf{r}_i + s\hat{\mathbf{n}}_i$, for $s \in [-l_i/2, l_i/2]$, along the cell's centerline, as described above; and $a_i(s)$ is the cell-substratum contact area density, which is the integrand in Eqn. 3:

$$a_i(s) = R^{1/2}\left(1 - \left|\hat{n}_{i,z}\right|^2\right)\delta_i^{1/2}(s) + \pi R\left|\hat{n}_{i,z}\right|^2 \Theta\left(\delta_i(s)\right).$$

To impose differential cell-substratum friction in the 3D ABMs, we assumed that:

1. the cell-substratum adhesion energy density, $\Sigma_{0,i}$, of each cell switches between $\Sigma_{0,\mathrm{H}}$ and $\Sigma_{0,\mathrm{L}}$ (Eqn. 3); and
2. the cell-substratum friction coefficient, $\eta_{1,i}$, of each cell switches between $\eta_{1,\mathrm{H}}$ and $\eta_{1,\mathrm{L}}$.

Note that a larger value for $\Sigma_{0,i}$ causes a concomitant decrease in the cell-substratum adhesion energy (Eqn. 3), which causes the cell to lower its *z*-position and increase its contact area with the substratum (by increasing $a_i(s)$), thereby also increasing the magnitude of $\mathbf{F}_{i,\mathrm{csf}}$. We fixed $\Sigma_{0,\mathrm{H}}$ and $\eta_{1,\mathrm{H}}$ at values previously estimated[1] for the Δ*rbmA* rugose *V. cholerae* strain, $\Sigma_{0,\mathrm{H}} = 7.704 \cdot 10^{-6}\ \mathrm{N \cdot m^{-1}}$ and $\eta_{1,\mathrm{H}} = 2 \cdot 10^5\ \mathrm{Pa \cdot s}$, and set $\Sigma_{0,\mathrm{L}} = 10^{-5}\Sigma_{0,\mathrm{H}}$ and $\eta_{1,\mathrm{L}} = 10^{-5}\eta_{1,\mathrm{H}}$.

***Differential cell-gel friction in 3D (Figures 5H, S6D, and S6E).*** We imposed differential cell-gel friction in the 3D ABMs by assuming that:

1. the cell-gel adhesion strength, $\Sigma_{\mathrm{gel},i}$, of each cell switches between $\Sigma_{\mathrm{gel,H}}$ and $\Sigma_{\mathrm{gel,L}}$ (see ref. [3] for details); and
2. the cell-gel friction coefficient, $\eta_{\mathrm{cg},i}$, of each cell switches between $\eta_{\mathrm{cg,H}}$ and $\eta_{\mathrm{cg,L}}$ (Eqn. 2).

The reasoning here is analogous to our scheme for imposing differential cell-substratum friction in 3D: the cell-gel friction force on cell $i$ due to gel particle $j$ (Eqn. 2) scales with the theoretical overlap, $\delta_{i,j}$, between the two bodies; in turn, this overlap scales with the cell-gel adhesion strength, $\Sigma_{\mathrm{gel},i}$ (see ref. [3] for details). We fixed $\Sigma_{\mathrm{gel,H}}$ at the value previously estimated[3] for the Δ*rbmA* rugose *V. cholerae* strain, $\Sigma_{\mathrm{gel,H}} = 6 \cdot 10^{-2}\ \mathrm{N \cdot m^{-2}}$, and set $\Sigma_{\mathrm{gel,L}} = 10^{-5}\Sigma_{\mathrm{gel,H}}$. While we did not have a direct experimental estimate for $\eta_{\mathrm{cg,H}}$, we assumed for simplicity that $\eta_{\mathrm{cg,H}} = \eta_{1,\mathrm{H}} = 2 \cdot 10^5\ \mathrm{Pa \cdot s}$, and similarly set $\eta_{\mathrm{cg,L}} = 10^{-5}\eta_{\mathrm{cg,H}}$.

***Differential ambient viscosity in 3D (Figure 5K).*** Finally, we imposed differential ambient viscosity in the 3D ABMs by assuming that the dynamic viscosity, $\eta_{0,i}$, of each cell switches between two values, $\eta_{0,\mathrm{H}}$ and $\eta_{0,\mathrm{L}}$ (see ref. [1] for details). Here, we fixed the *low*-c-di-GMP value, $\eta_{0,\mathrm{L}}$, at the value previously estimated[1] for the Δ*rbmA* rugose *V. cholerae* strain, $\eta_{0,\mathrm{L}} = 20\ \mathrm{Pa \cdot s}$ (which is roughly $0.072\ \mathrm{kg \cdot \mu m^{-1} \cdot hr^{-1}}$), and set $\eta_{0,\mathrm{H}} = 10^4\eta_{0,\mathrm{L}}$.

**4. Growth void in 2D ABMs.**

We also implemented a collection of 2D ABMs with a central growth void, as a proxy for the verticalized core[2] (Figure 5G). In each of these simulations, a growth void with a particular normalized radius, $\rho_{\text{void}} \in [0,1]$, was introduced at each timepoint in such a way that, for each azimuthal angle, the innermost $\rho_{\text{void}}$ fraction of cells along the corresponding radial direction from the origin were assigned a zero growth rate for all subsequent time. In practice, this was done at each timepoint as follows:

1. Determine the origin of the biofilm as the mean over all cell centers (i.e., the center of mass).
2. Collect, for each cell, five equally spaced points from one endpoint of the centerline to the other.
3. Calculate an $\alpha$-shape[15] of these points (see below), and include each cell that contributes a point to the $\alpha$-shape as part of the biofilm's periphery.
4. For each non-peripheral cell, with radial distance $r$ and azimuthal angle $\theta$ with respect to the origin:
   a. Find the peripheral cell with the greatest radial distance from the origin among those whose azimuthal angle is within $5°$ of $\theta$; if no such peripheral cell exists, find the peripheral cell with the azimuthal angle closest to $\theta$. Let the radial distance of the chosen peripheral cell be $r_{\text{max}}$.
   b. If $r/r_{\text{max}} < \rho_{\text{void}}$, then set the growth rate of the non-peripheral cell to zero.

The value of $\alpha$ in the $\alpha$-shape, which controls the granularity of the $\alpha$-shape[15,16], was chosen such that the $\alpha$-shape is as granular as possible while being simply connected (i.e., the points in the $\alpha$-shape form one contiguous cycle that encloses all the other points).

To compare distributions of cell velocities in simulations with and without the growth void (Figure 5G), we calculated the velocity of each cell $i$ in each simulation as the forward difference,

$$\mathbf{v}_i(t) = \frac{\mathbf{r}_i(t+\Delta t) - \mathbf{r}_i(t)}{\Delta t},$$

with $\Delta t$ set to 3 min. The radial velocity vector was then calculated as the projection of $\mathbf{v}_i(t)$ onto the radial direction,

$$\mathbf{v}_{r,i}(t) = \left(\mathbf{v}_i(t) \cdot \frac{\mathbf{r}_i(t) - \langle \mathbf{r}_i(t) \rangle}{\|\mathbf{r}_i(t) - \langle \mathbf{r}_i(t) \rangle\|}\right)\left(\frac{\mathbf{r}_i(t) - \langle \mathbf{r}_i(t) \rangle}{\|\mathbf{r}_i(t) - \langle \mathbf{r}_i(t) \rangle\|}\right),$$

where the average, $\langle \mathbf{r}_i(t) \rangle$, was taken over all cells at time $t$. The "radial velocity" we refer to in the main text is the signed magnitude of this vector, $\epsilon_i \|\mathbf{v}_{r,i}(t)\|$, where $\epsilon_i = +1$ if $\mathbf{v}_{r,i}(t)$ points in the same direction as $\mathbf{r}_i(t) - \langle \mathbf{r}_i(t) \rangle$ and $\epsilon_i = -1$ otherwise.

To calculate the distributions in Figure 5G, we ran 20 simulations with the growth void ($\rho_{\text{void}} = 0.5$) and without, with mean state lifetimes of $\tau_{\text{L}} = \tau_{\text{H}} = 8\ \text{h}$ and with differential drag imposed via cell-substratum friction coefficients of $\eta_{1,\text{H}} = 720\ \text{kg} \cdot \mu\text{m}^{-1} \cdot \text{hr}^{-1}$ and $\eta_{1,\text{L}} = \eta_{1,\text{H}}/20$ (see **Section 3** and **Table 1**). (Here, we set $\tau_{\text{L}} = \tau_{\text{H}} = 8\ \text{h}$, instead of the experimentally estimated values of $\tau_{\text{L}} = 2\ \text{h}$ and $\tau_{\text{H}} = 4\ \text{h}$, to examine the effects of the growth void in the 2D scenario where sorting is most pronounced, as shown in Figure S6B.) We quantified the radial velocity of each cell in each simulation at time $t = t_{\text{max}} - \Delta t$, where $t_{\text{max}}$ is the timepoint by which the biofilm first reached 5000 cells in each simulation. Shown in the *bottom* violin in Figure 5G are the radial velocity distributions of the low-c-di-GMP (red) and high-c-di-GMP (cyan) cells whose normalized radial distances,

$$\frac{\|\mathbf{r}_i(t) - \langle \mathbf{r}_i(t) \rangle\|}{\max_i\{\|\mathbf{r}_i(t) - \langle \mathbf{r}_i(t) \rangle\|\}},$$

are between 0.55 and 0.75, across 20 simulations *with* the growth void. These are the cells whose radial alignment due to the growth void—whose radius was fixed at $\rho_{\text{void}} = 0.5$—is strongest[2]. In contrast, shown in the *top* violin in Figure 5G are the corresponding distributions for cells whose radial distances are between 0.05 and 0.25, across 20 simulations *without* the growth void. This group of cells was chosen to match the strength of the growth-induced flow experienced by the two groups of cells.

**5. Implementational details.** The 3D ABM was implemented in LAMMPS[18], as described previously[2–4]. The 2D ABM was implemented in C++, using the Boost[19] (version 1.83.0), Eigen[20] (version 3.4.0), and CGAL[21] (version 5.6) libraries as dependencies. An adaptive Runge–Kutta method, due to Bogacki and Shampine, was used for time-integration of the equations of motion[22]; timesteps were adapted according to the Runge–Kutta error estimate as prescribed in ref. [23]. For growth void simulations, $\alpha$-shapes were computed using a custom C++ routine that adapted the CGAL class `Alpha_shape_2`.

| Parameter | Meaning | Default value |
|---|---|---|
| $R$ | Cell radius, including matrix coating | $0.8\ \mu\text{m}$ |
| $R_{\text{cell}}$ | Cell body radius | $0.5\ \mu\text{m}$ |
| $\gamma_0$ | Mean growth rate (s.d. = $0.2\gamma_0$) | $1.12\ \text{hr}^{-1}$ |
| $l_0$ | Cell length at birth | $1\ \mu\text{m}$ |
| $E_0$ | Elastic modulus of matrix coating | $3900\ \text{kg} \cdot \mu\text{m}^{-1} \cdot \text{hr}^{-2}$ |
| $E_{\text{cell}}$ | Elastic modulus of cell body | $10^3 E_0$ |
| $\Sigma_0$ | Cell-substratum adhesion energy density | $100\ \text{kg} \cdot \text{hr}^{-2}$ |
| $\eta_0$ | Dynamic viscosity due to matrix | $0.072\ \text{kg} \cdot \mu\text{m}^{-1} \cdot \text{hr}^{-1}$ |
| $\eta_1$ | Cell-substratum friction coefficient | $720\ \text{kg} \cdot \mu\text{m}^{-1} \cdot \text{hr}^{-1}$ |

**Table 1: Default parameter values for the 2D ABMs.** The values given here are roughly equivalent to their corresponding values in the 3D ABMs (**Table 2**).

| Parameter | Meaning | Default value |
|---|---|---|
| $R$ | Cell radius, including matrix coating | $0.8\ \mu\mathrm{m}$ |
| $R_{\mathrm{cell}}$ | Cell body radius | $0.5\ \mu\mathrm{m}$ |
| $\gamma_0$ | Mean growth rate (s.d. = $0.2\gamma_0$) | $3.12 \cdot 10^{-4}\ \mathrm{s}^{-1}$ |
| $l_0$ | Cell length at birth | $1\ \mu\mathrm{m}$ |
| $E_0$ | Elastic modulus of matrix coating | $300\ \mathrm{Pa}$ |
| $E_{\mathrm{cell}}$ | Elastic modulus of cell body | $10^3 E_0$ |
| $\Sigma_0$ | Cell-substratum adhesion energy density | $7.704 \cdot 10^{-6}\ \mathrm{N} \cdot \mathrm{m}^{-1}$ |
| $\eta_0$ | Dynamic viscosity due to matrix | $20\ \mathrm{Pa} \cdot \mathrm{s}$ |
| $\eta_1$ | Cell-substratum friction coefficient | $2 \cdot 10^5\ \mathrm{Pa} \cdot \mathrm{s}$ |
| $R_{\mathrm{gel}}$ | Gel particle radius | $0.8\ \mu\mathrm{m}$ |
| $E_{\mathrm{gel}}$ | Gel elastic modulus | $10^4\ \mathrm{Pa}$ |
| $k_r$ | Gel inter-particle spring constant | $8 \cdot 10^{-3}\ \mathrm{N} \cdot \mathrm{m}^{-1}$ |
| $\xi_0$ | Equilibrium gel inter-particle distance | $0.8\ \mu\mathrm{m}$ |
| $\Sigma_{\mathrm{gel}}$ | Cell-gel adhesion strength | $6 \cdot 10^{-2}\ \mathrm{N} \cdot \mathrm{m}^{-2}$ |
| $\eta_{\mathrm{cg}}$ | Cell-gel friction coefficient | $2 \cdot 10^5\ \mathrm{Pa} \cdot \mathrm{s}$ |

**Table 2: Default parameter values for the 3D ABMs.** The gel parameters were calibrated to mimic a 1.5% agarose gel.

| Parameter | Meaning | Value |
|---|---|---|
| $R_{\mathrm{gel}}$ | Gel particle radius | $0.8\ \mu\mathrm{m}$ |
| $E_{\mathrm{gel}}$ | Gel elastic modulus | $333\ \mathrm{Pa}$ |
| $k_r$ | Gel inter-particle spring constant | $2.4 \cdot 10^{-4}\ \mathrm{N} \cdot \mathrm{m}^{-1}$ |
| $\xi_0$ | Equilibrium gel inter-particle distance | $0.8\ \mu\mathrm{m}$ |
| $\Sigma_{\mathrm{gel}}$ | Cell-gel adhesion strength | $0$ |
| $\eta_{\mathrm{cg}}$ | Cell-gel friction coefficient | $0$ |

**Table 3: Soft gel parameter values for the 3D ABMs.** These gel parameters were calibrated to mimic a 0.225% agarose gel.

**References.**


1. Beroz, F., Yan, J., Meir, Y., Sabass, B., Stone, H.A., Bassler, B.L., and Wingreen, N.S. (2018). Verticalization of bacterial biofilms. Nat. Phys. *14*, 954–960. https://doi.org/10.1038/s41567-018-0170-4.

2. Nijjer, J., Li, C., Zhang, Q., Lu, H., Zhang, S., and Yan, J. (2021). Mechanical forces drive a reorientation cascade leading to biofilm self-patterning. Nat. Commun. *12*, 6632. https://doi.org/10.1038/s41467-021-26869-6.

3. Nijjer, J., Li, C., Kothari, M., Henzel, T., Zhang, Q., Tai, J.-S.B., Zhou, S., Cohen, T., Zhang, S., and Yan, J. (2023). Biofilms as self-shaping growing nematics. Nat. Phys. *19*, 1936–1944. https://doi.org/10.1038/s41567-023-02221-1.

4. Li, C., Nijjer, J., Feng, L., Zhang, Q., Yan, J., and Zhang, S. (2024). Agent-based modeling of stress anisotropy driven nematic ordering in growing biofilms. Soft Matter *20*, 3401. https://doi.org/10.1039/d3sm01535a.

5. Landau, L.D., and Lifshitz, E.M. (1986). Theory of Elasticity 3rd ed. (Butterworth-Heinemann).

13. Norris, J.R. (1997). Markov Chains (Cambridge University Press).

19. Abrahams, D., and others (2023). Boost. Version 1.83.0.

20. Guennebaud, G., Jacob, B., and others (2021). Eigen. Version 3.4.0.

21. The CGAL Project (2023). CGAL. Version 5.6.

22. Bogacki, P., and Shampine, L.F. (1989). A 3(2) pair of Runge–Kutta formulas. Appl. Math. Lett. *2*, 321–325. https://doi.org/10.1016/0893-9659(89)90079-7.

23. Press, W.H., Teukolsky, S.A., Vetterling, W.T., and Flannery, B.P. (2007). Numerical Recipes: the Art of Scientific Computing 3rd ed. (Cambridge University Press).
